# UPP affects chloroplast development by interfering with chloroplast proteostasis

**DOI:** 10.1101/2023.08.28.555145

**Authors:** Vanessa Scherer, Leo Bellin, Serena Schwenkert, Martin Lehmann, Jannis Rinne, Claus-Peter Witte, Kathrin Jahnke, Andreas Richter, Tobias Pruss, Anne Lau, Lisa Waller, Sebastian Stein, Dario Leister, Torsten Möhlmann

## Abstract

Arabidopsis uracil phosphoribosyltransferase (UPP) is an essential enzyme which appears to have a previously unknown, moonlighting activity. Our analysis of UPP *amiRNA* mutants has confirmed that this vital function is crucial for chloroplast development and growth. Interestingly, this function appears to be unrelated to nucleotide homeostasis since nucleotide levels were not altered in the studied mutants. Transcriptomics and proteomic analysis suggest that UPP plays a role in chloroplast proteostasis, especially under high light (HL). Immunoblots of mature plants and a de-etiolation experiment with young seedlings revealed PetC, the iron-sulfur protein of the cytochrome *b*_6_*f* complex, as a putative UPP target. In addition, UPP and PetC were identified in a high molecular weight complex. Consistently, we show that PetC is massively reduced in UPP knock-down plants.

The block observed in photosynthetic electron transport, as evidenced by reduced high light-induced non-photochemical quenching (NPQ) but increased unregulated energy dissipation (NO), might be therefore a consequence of reduced PetC. After HL treatment, UPP *amiRNA* mutants showed impaired photosynthesis and reduced carbohydrate contents, resulting in an inability to induce flavonoid biosynthesis. In addition, the levels of the osmoprotectants raffinose, proline and fumarate were found to be reduced. Proteases, including thylakoid filamentation temperature-sensitive 1, 5 (FtsH), caseinolytic protease proteolytic subunit 1 (ClpP1), and processing peptidases, as well as components of the chloroplast protein import machinery, were up-regulated.

In sum, our work suggests that UPP assists in stabilizing and protection of PetC during the assembly of the Fe-S cluster and targeting to the thylakoid.

## Introduction

Nucleotides are organic molecules that are essential components of genetic material and play important roles in cellular processes. Uracil phosphoribosyl transferase (UPP) catalyses the first committed step in uracil salvage and is located in the chloroplast. Arabidopsis recombinant UPP converts uracil and phosphoribosyl pyrophosphate into uridine monophosphate (UMP). UPP knockout mutants (*upp-1*) do not exhibit uracil phosphoribosyl transferase (UPRT) activity, indicating that no other protein exhibits UPRT activity (Mainguet et al., 2009; Ohler et al., 2019). The functionality of uracil salvage in vivo was demonstrated by 5-fluorouracil toxicity and consequential resistance of mutants with reduced UPRT activity (Ohler et al., 2019). Seedling lethal upp-1 can be rescued by complementation with nearly catalytic inactive UPP but not with a highly active homolog from *Escherichia coli*. This agrees with previous results and suggests that an unknown second (moonlighting) function is responsible for the inability of *upp-1* to establish photosynthesis (Mainguet et al., 2009; Arrivault, 2019; Ohler et al., 2019).

Photosynthesis is a dominant process in green plant tissues. Chloroplasts have their own machinery for transcription and translation of several photosynthesis genes. Mutants in nucleotide metabolism that exhibit nucleotide limitation also affect photosynthesis. Examples of knock-down mutants targeting enzymes involved in pyrimidine de novo synthesis include aspartate transcarbamoylase (ATC), which catalyses the first committed step, CTP-synthase 2 (CTPS2), which catalyses the final step, as well as thymidine kinase 1b (TK1b) and double knock-out mutants for uridine-cytidine kinase (UCK) 1 and 2 (Clausen et al., 2012; Ohler et al., 2019; Bellin et al., 2021a; Bellin et al., 2021b). The rescue of ATC and CTPS mutants by nucleotide precursors provides evidence that nucleotide limitation is the cause of the observed effects on photosynthesis.

Previous work has provided evidence for a moonlighting function of Arabidopsis UPP in chloroplast development and the establishment of photosynthesis, as reported by Ohler et al. (2019). Additionally, downstream enzymes such as plastid UMP kinase (PUMPKIN) and nucleoside diphosphate kinase 2 (NDPK2) have also been found to exhibit moonlighting functions (Dorion and Rivoal, 2015; Arrivault, 2019; Schmid et al., 2019). Since knock-out mutants of both enzymes are not lethal to seedlings, as is the case with UPP, these moonlighting functions are likely distinct from those of UPP.

Chloroplast differentiation, which leads to photosynthesizing chloroplasts, occurs in two distinct phases: the Structure Establishment Phase and the Chloroplast Proliferation Phase (Pipitone et al., 2021). The establishment of the structure phase is associated with the formation of thylakoid membranes and the assembly of the photosystems (PSII, PSI, cytochrome *b*_6_*f* (Cyt *b*_6_*f*)) within the first 24 hours after de-etiolation (Pipitone et al., 2021). However, the biogenesis and assembly of the photosystems, particularly cytb6/f, remain poorly understood. A new assembly factor for this complex was recently identified as de-etiolation induced protein 1 (DEIP1). DEIP1 is a 25 kDa protein located in the thylakoid that interacts with PetA and PetB from cytb6/f during assembly (Sandoval-Ibanez et al., 2022).

This work proposes a function of UPP in chloroplast proteostasis, primarily by affecting PetC stability and assembly into cytb6/f. This hypothesis is supported by transcriptomics and proteomics of UPP-*amiRNA* lines (*ami-upp*). Additionally, immunoblotting of mature and de-etiolated Arabidopsis plants provides further evidence. Reduced PetC amounts are accompanied by increased levels of chloroplast protein import machinery components and proteases, such as thylakoid filamentation temperature-sensitive (FtsH) metalloproteases involved in the assembly or repair of PSII, PSI, and Cyt *b*_6_*f* complexes (Malnoe et al., 2014; Jarvi et al., 2016; Kato et al., 2018).

## Results

### Although nucleotide levels are wild type like, UPP *amiRNA* lines exhibit reduced growth

To generate UPP *amiRNA* lines with reduced amounts of UPP transcript, three lines were used for further analysis: *ami-upp-1*, *ami-upp-4*, and *ami-upp-7*, with 54%, 34%, and 4.6% of wild type (Col-0) transcript levels. The reductions in fresh weight of these mutants after six weeks of growth on soil correlated well with the lowered amounts of UPP protein and UPP transcript (Figure 1A-C). The protein levels were quantified using a specific antibody (Ohler et al., 2019). The results showed reduced UPP amounts of 97%, 7%, 13%, 11%, 2%, and 2% compared to Col-0 for *ami-upp-1*, -*3*, -*4*, - *5*, -*6*, and *ami-upp-7*, respectively (Figure 1C). This study will focus on *ami-upp-1*, *ami-upp-4*, *ami-upp-7*, and a complementation line expressing UPP under control of the ubiquitin 10 promoter in *upp-1* (knock-out) background (*UBQ10::UPP*, Ohler et al., 2019), referred to as *UPP*-OX.

**Figure 1.**
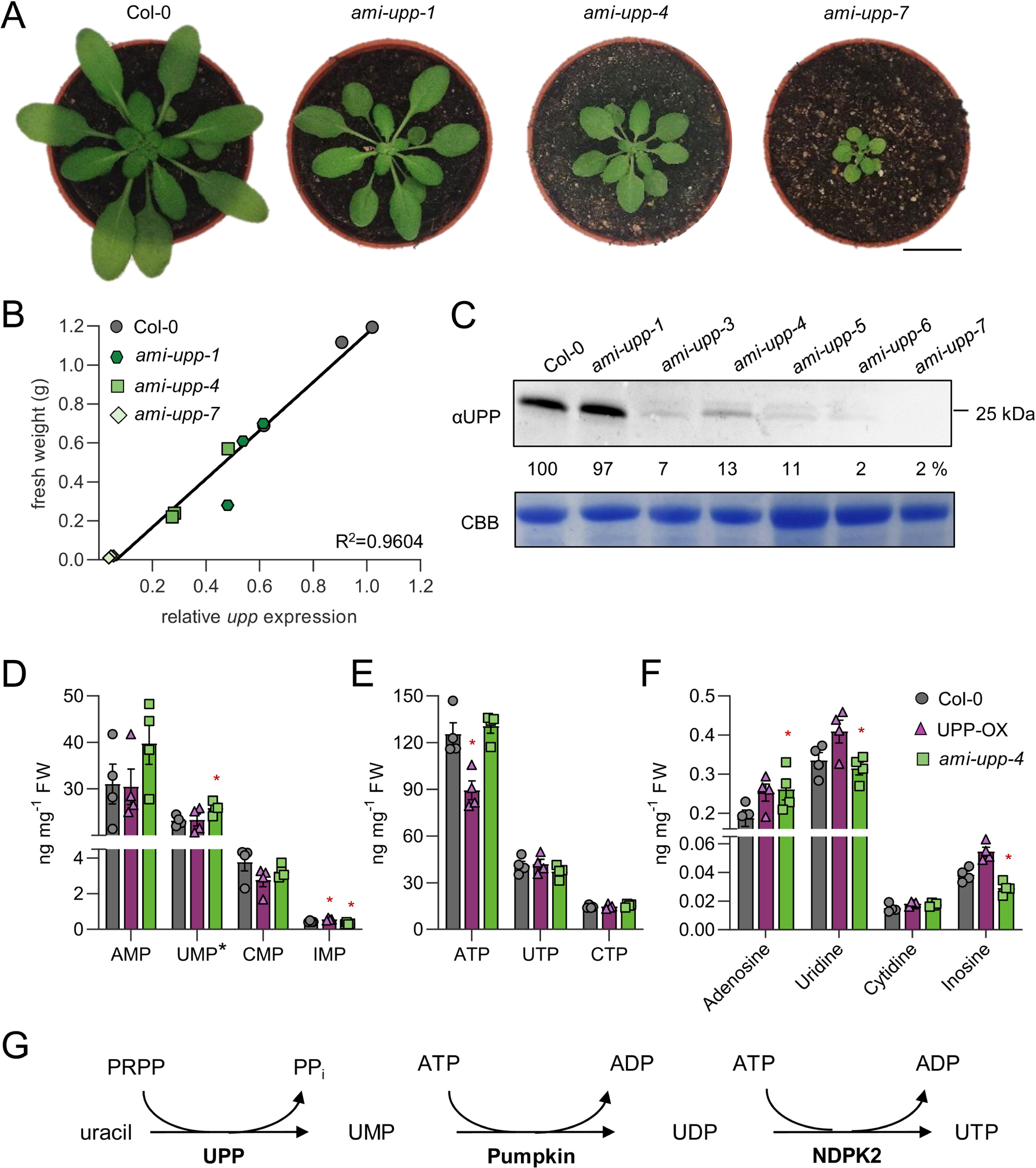
Reduced UPP contents provoke severely impaired plant growth and development. (**A**) Representative five-week old Col-0 and *UPP* knock-down plants (*ami-upp*) grown under a short day regime on soil (10h light/ 14h dark). (**B**) Correlation of relative *upp* expression and fresh weight of five-week old plants (n = 3). (**C**) Western blot detection of UPP amounts with a corresponding quantification in percent to Col-0. Coomassie-brilliant blue (CBB) staining as loading control. (**D**) Nucleotide-mono-phosphates, UMP*, values for UMP are the sum of free UMP and some UMP released from a low degree of UDP-glucose decay during sample preparation (**E**) nucleotide-triphosphates and (**F**) nucleoside concentrations in five-week old plants (n = 4). (**G**) Scheme of the uracil salvage pathway in the chloroplast. Plotted are the mean values of at least three biological replicates with the corresponding standard deviation. For statistical analysis one-way ANOVA (**D**, **E**, **F**) was performed followed by Dunnetts’s multiple comparison test (* = p < 0.05; ** = p < 0.01; *** = p < 0.001). Scale bar in **A** = 2 cm. pumpkin, plastidic UMP kinase; NDPK2, nucleoside diphosphate kinase 2 (plastid).

UPP is an enzyme involved in pyrimidine salvage and has a secondary function that affects the development of UPP mutants (Ohler et al., 2019). To test this finding, we quantified levels of (deoxy)nucleotides, nucleosides, and nucleobases in *ami-upp-4* and *UPP*-OX. There were no significant differences in the levels of these molecules between the two genotypes (Figure 1D, Supplemental Figure S1, 2). It is unlikely that UPP plays a direct role in nucleotide metabolism that is fundamental for the distinct growth phenotypes of *amiRNA* mutants. The analysis included plants of all genotypes grown at 4°C or in HL (high light, 1000 µmol photons s^−1^ m^−2^). Comparing between conditions revealed some alterations, such as increased levels of purine nucleotides ATP, ADP and dADP in one day cold-treated plants (see Supplementary Figure S1). In HL-treated plants, both ADP and dADP increased, whereas UDP and GDP decreased (see Supplemental Figures S1 and S2).

### Transcriptome analysis points to alterations in the functional category “protein”

RNA-Seq analysis of four-week-old UPP mutants grown under control conditions or under high light for 3 hours revealed a high number of differentially expressed genes (DEGs). We included the high light treatment because we had already observed a high light-dependent lack of anthocyanin accumulation in *ami-upp* lines (see below). Under <u>m</u>edium light conditions (ML, 120 µmol light), 252 differentially expressed genes (DEGs) were found in *ami-upp-4* compared to Col-0, with a padj. value of < 0.05. However, after high light (HL, 1000 µmol) treatment, this number increased to 2502 DEGs in the same comparison together with a higher scattering of expression values when including all genes which were identified (Figure 2A-C). Among genes encoding non plastid proteins (padj. < 0.05 and log_2_-fold change <−1 or >1) qua-quine starch (QQS) and cytosolic ribosomal protein P2D (RPP2D) were found downregulated in ML and HL conditions. QQS represents a regulator of carbon and nitrogen allocation to starch and protein (Li and Wurtele, 2015). When focusing on the expression of nuclear genes encoding chloroplast proteins (log fold change <−1 or >1), only three altered genes were identified in ML-grown plants and five genes in HL-grown plants. The upregulation of both early light-induced proteins (ELIP) 1 and 2 was more pronounced under high light conditions compared to moderate light conditions (Figure 2D, E). ELIPs function as regulators of chlorophyll synthesis and protect against light-induced damage by binding free chlorophyll molecules (Tzvetkova-Chevolleau et al., 2007). Additionally, high expression levels of ELIPs are observed 4 to 24 hours after de-etiolation (Pipitone et al., 2021). An analysis of GO terms for significantly differentially expressed genes (DEGs) in HL-grown plants revealed a high abundance of genes related to the categories of ‘protein’ and ‘ribosomal proteins’, while the categories of ‘stress’ and ‘abiotic stress’ were downregulated (Figure 2F).

**Figure 2.**
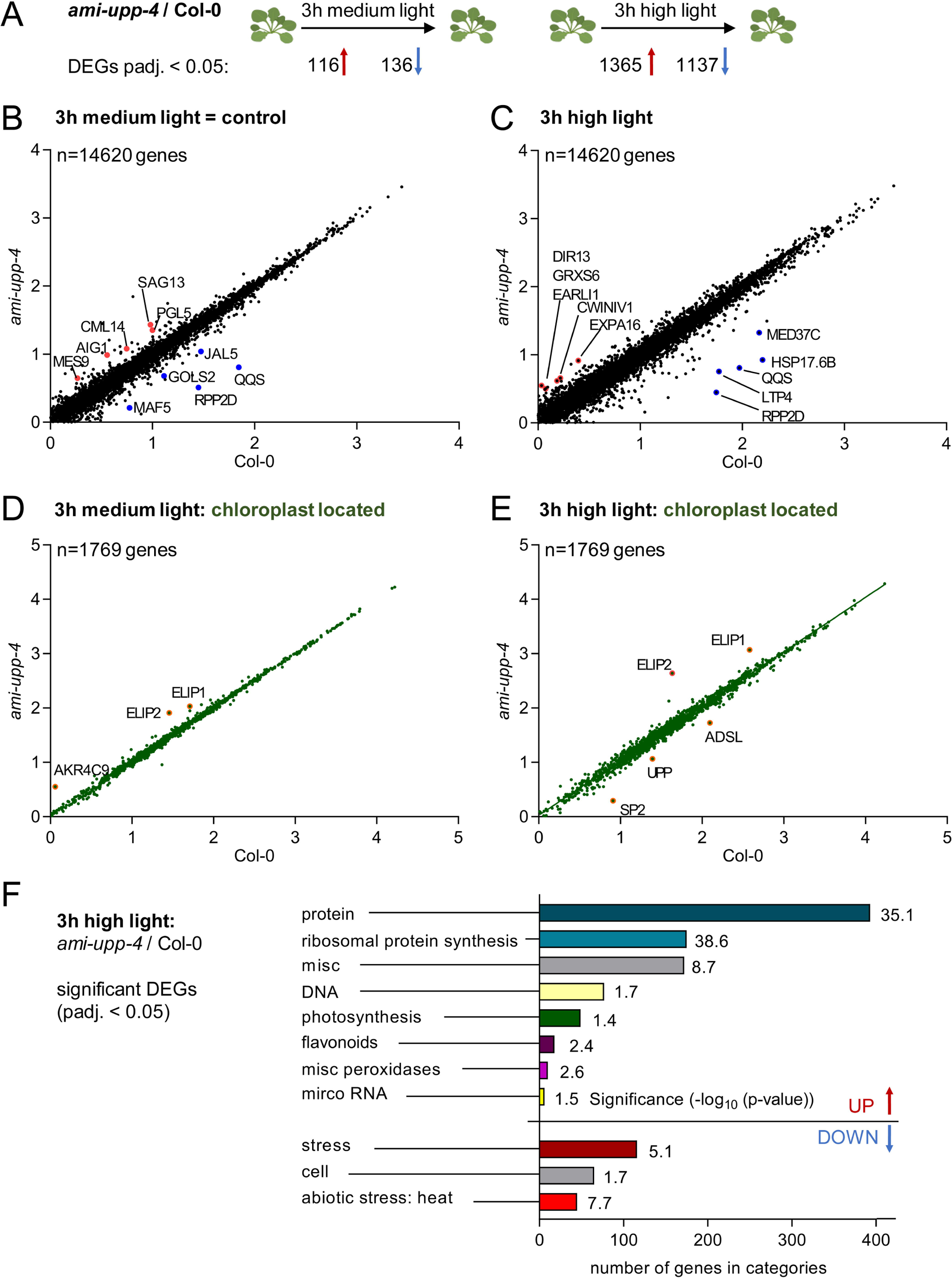
Gene expression analysis showed alterations in *ami-upp-4* under high light stress. (**A**) RNA Seq analysis of different expressed genes (DEGs) in *ami-upp-4* compared to Col-0 under three hours of medium light (= control) and three hours of high light (1000 μmol photons m^−2^ s^−1^). Plants were grown for five weeks grown under a short-day regime on soil (10h light/ 14h dark). Scattered plot of all up- and down-regulated genes found in *ami-upp-4* and Col-0 under control (**B**) and high light (**C**) conditions. Three biological replicates were analyzed per time point. All genes located in the chloroplast, from both control (**D**) and high light (**E**) conditions, were plotted in a separate scatter plot. **B-E** Labeled genes filtered by padj. < 0.05, log_2_ fold change >1, <-1, all units: log_10_ FPKM. (**F**) GO-term analysis of significantly responded Mapman bins after Benjamini-Hochberg correction (*padj.* < 0.05).

### Proteomics reveal distinct alterations in proteostasis especially upon HL treatment

Label-free proteomics was selected as an objective method to investigate the moonlighting function of UPP. Whole rosette material from five-week-old plants was exposed to HL treatment after two hours of illumination with ML (0h HL). Three hours of HL (3h HL), eight hours of HL (8h HL), and subsequent 12 hours of dark recovery followed by 2h HL (recovery) were also tested. Samples from all time points were processed in three independent biological replicates. The untreated samples exhibited 104 significantly (p-value < 0.05 in pairwise comparison) upregulated and 113 downregulated proteins in *ami-upp-4* compared to Col-0 (Figure 3A, B). This number increased to 151 upregulated and 209 downregulated proteins after 3h HL, of which 23% were located in the chloroplast (Figure 3A). The most significant increase was observed after 8h HL, with 380 upregulated and 303 downregulated proteins, of which 27% were located in the chloroplast (Figure 3A, C). After recovery, 166 proteins were still upregulated and 245 downregulated compared to Col-0 (Figure 3A). The 8h HL proteome underwent a GO-term analysis using the Mapman tool (Thimm et al., 2004). The analysis showed significant changes in categories 29 (protein), 20 (stress), 21 (redox), and 16 (secondary metabolism) (Figure 3D). Bin 29 (protein) comprises 158 proteins, of which 48% are differentially expressed proteins (DEPs) involved in protein-synthesis, followed by 23% in protein-degradation, 10% in amino acid activation, 8% in protein-targeting, 4.5% in posttranslational modification, 4.5% in protein-folding, and 2% in assembly and cofactor ligation (see Figure 3E).

**Figure 3.**
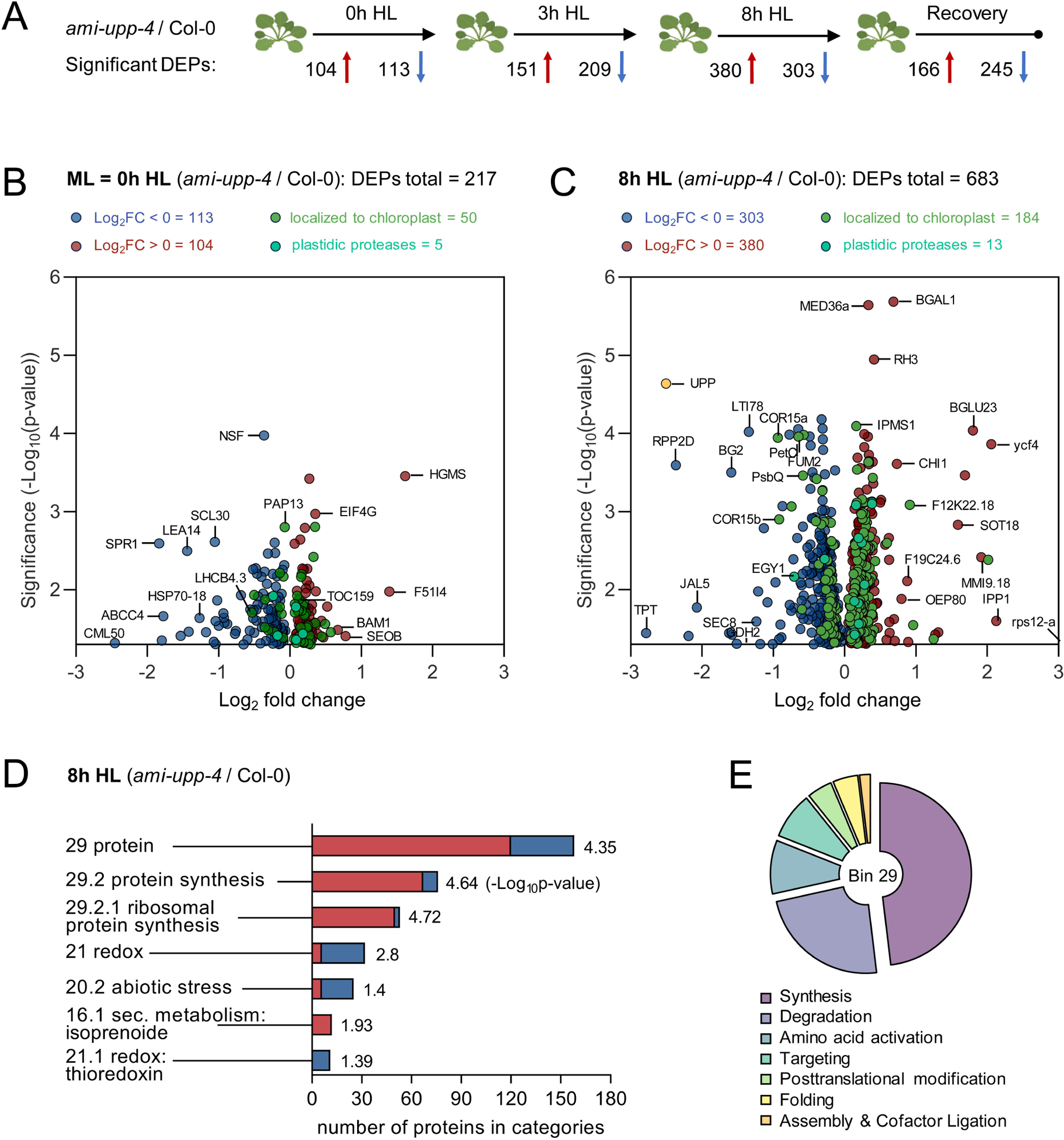
Proteomics reveal distinct alterations in proteostasis under high light stress. **(A)** Workflow of proteome experiment. HL (1000 μmol photons m^−2^ s^−1^) was applied for the indicated time period and differentially (p-value < 0.05) up and downregulated proteins were analyzed (numbers are indicated). Three biological replicates were analyzed per time point. DEPs were analyzed in pairwise comparison using t-test. (**B, C**) Volcano plots show DEPs (*p* < 0.05) in *ami-upp-4* compared to Col-0 under control (**B**) and after eight hours high light treatment (**C**). Proteins localized to the chloroplast are highlighted in dark green and plastidic proteases in bright green. (**D**) GO-term analysis of significantly responded Mapman bins after Benjamini-Hochberg correction (*padj.* < 0.05) and (**E**) further subdivision of bin 29, red = upregulated proteins, blue = down-regulated proteins. (* = *p* < 0.05; ** = *p* < 0.01; *** = *p* < 0.001).

When focusing proteins related to photosynthesis or proteostasis, we observed that rps12-A (chloroplast ribosomal protein), IPP1 (isopentenyl diphosphate:dimethylallyl diphosphate isomerase), YCF4 (photosystem assembly factor), and OEP80 (outer envelope protein 80) were upregulated (see Figure 2C). In contrast, the following genes/proteins were down-regulated after 8h HL: triosephosphate-phosphate-translocator (TPT), UPP, RPP2D, mitochondrial glycine cleavage system H-protein (GDH2), Rieske FE-S protein (PetC), small PSII protein (PsbQ2), and cytosolic fumarate hydratase involved in cold and high light acclimation (FUM2) (Figure 3C).

The protein groups that are markedly affected include those involved in electron transport reactions, chloroplast protein import and folding, and proteases (as shown in Figure 4A-C). Other groups consist of ribosomal proteins and redox processes, as well as the CBB (Calvin-Benson-Bassham cycle) (as depicted in Supplemental Figure S3). It was found that the light harvesting complexes, PSII subunits and Cytb6/f subunits were mostly reduced, whereas those of ATP-synthase, NDH (NAD(P)H dehydrogenase subunits involved in cyclic electron flow) and FNR (Ferredoxin NADP+ reductase) were increased (Figure 4A). Additionally, several components of chloroplast protein import were found to be altered. For instance, the proteins Toc33, Toc159, Toc75-3, and Tic110, which are part of the translocon of the outer and inner chloroplast envelope, respectively, were found to be increased (Figure 4B). Additionally, chaperones from the HSP and CPN60 families were observed to be altered at different time points in HL. OEP (outer envelope protein) 161 and 163 were reduced together with a factor (AKR2) mediating import of outer envelope membrane proteins (Bae et al., 2008).

**Figure 4.**
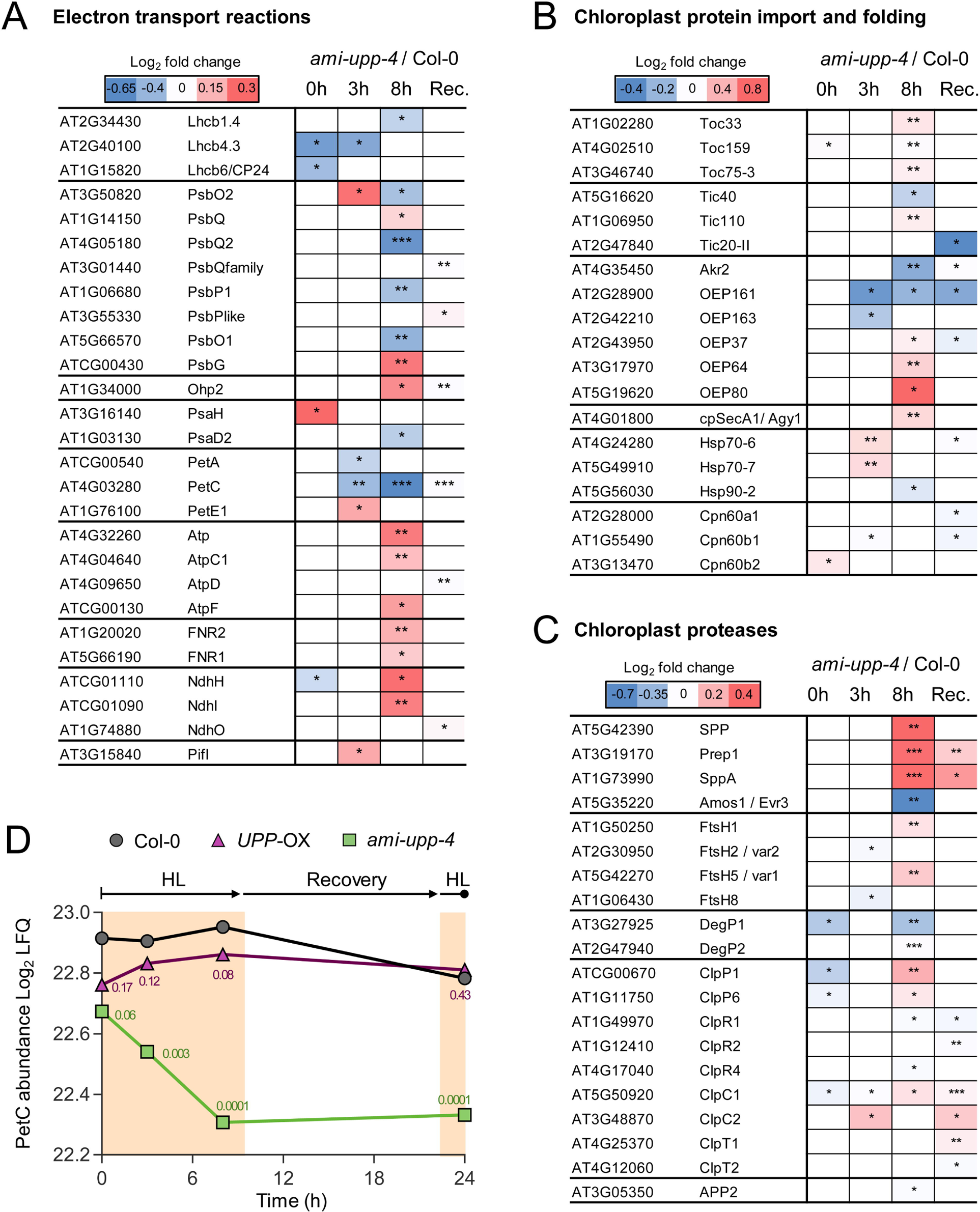
Proteome mass spec analysis reveals alterations in plastidic proteins in *ami-upp-4* mutants upon high light treatment. (**A**, **B**, **C**) Comparison of protein alterations between Col-0 and *ami-upp-4* associated to electron transport reactions (**A**), chloroplast protein import and folding (**B**) and chloroplast proteases (**C**) (*p* < 0.05). Plants were grown for five weeks grown under a short-day regime on soil (10h light/ 14h dark). (**D**) Proteome mass spec quantification (Log_2_ LFQ) of PetC from a HL time course experiment. P-values of a pairwise comparison to the corresponding Col-0 are shown. Plotted are the mean values of three biological replicates with the corresponding standard deviation. For statistical analysis *ami-upp-4* proteins were compared to Col-0 proteins in the same condition using Students t-test (* = *p* < 0.05; ** = *p* < 0.01; *** = *p* < 0.001).

In contrast, the amounts of OEP64 and OEP80 were increased after 8 hours of HL. Additionally, spSecA1, a motor protein involved in the cpSec1 pathway for ATP-driven import of nuclear-encoded proteins into thylakoids, was upregulated (Figure 4B).

Furthermore, several chloroplast proteases were found to be altered. After 8 hours of HL exposure, stromal and thylakoid processing peptidases SPP, SPPA, and Prep1 (presequence peptidase 1) increased, similar to thylakoid-bound FtsH (filamentation temperature sensitive-H) 1 and 5 (=variegated1, var1) (Figure 4C). DEGP1 (degradation of periplasmic proteins 1) and 2 decreased, whereas members of the Clp (caseinolytic protease) family mostly increased after 8 hours of HL or recovery. A metalloprotease is necessary for grana development and works together with FtsH2 (var2) to regulate chloroplast development. Its expression is significantly reduced after 8 hours of high light exposure (Figure 4C).

### PetC levels are reduced and thylakoid located FtsH proteases are up in *ami-upp* mutants

As many changes in the chloroplast proteome were detected, most prominent for PetC (Figure 4D) the steady-state levels of selected photosynthesis-related proteins were quantified through immunoblotting. The most significant change in abundance was again observed with PetC, which was greatly reduced in abundance in *ami-upp-4* and *-7* under both control and 8h HL (Figure 5A). An upregulation of FtsH 1/5 was observed, most notably in HL treated *ami-upp-7*. Both *ami-upp* mutants showed upregulation of PsbS under ML and HL conditions, while all other tested proteins remained nearly unchanged (Figure 5A). PsbS, also known as NPQ4, activates non-photochemical quenching (NPQ) and regulates high light acclimation by enhancing reactive oxygen species (ROS) homeostasis (Correa-Galvis et al., 2016). Additionally, proteomics analysis revealed an increase in FtsH1 and FtsH5 after 8h HL and recovery (Supplemental Figure S4). We conducted a de-etiolation experiment to determine if impaired PetC accumulation was observed shortly after illumination. After 7 days of growth in complete darkness, we observed only low accumulation of UPP and PetC. However, after only 24 hours of illumination, PetC levels were strongly induced together with UPP in Col-0, but not in *petc-2* (Figure 5B). The amount of other tested proteins was hardly affected.

**Figure 5.**
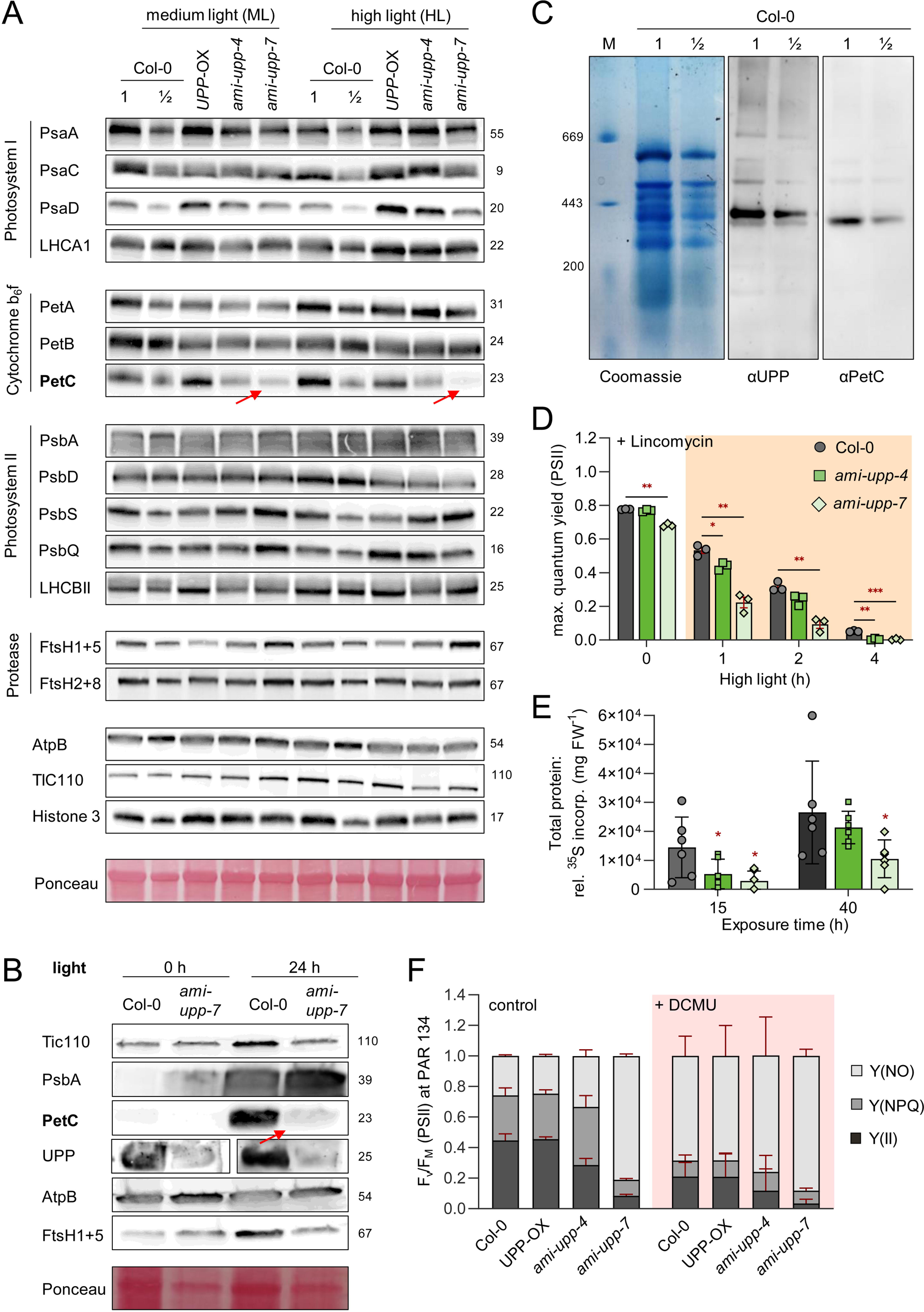
PetC protein is reduced and proteostasis is disturbed in *ami-upp* mutants. (**A**) Immunoblot of photosynthesis proteins in leaves of 5-week-old plants after 2 h of medium light (ML; 120) or additionally 8 h of high light (HL; 1000 μmol photons m^−2^ s^−1^) with corresponding ponceau staining as loading control. (**B**) Immunoblot of chloroplast located proteins in de-etiolated seedlings of Col-0 and *ami-upp-7* after 2 h light followed by 7 days in the dark and additional 24 h in the light. As loading control, a ponceau staining was used. (**C**) Blue Native-polyacrylamide gel electrophoresis of wildtype thylakoid membranes, stained with Coomassie-brilliant blue, blotted on PVDF membranes and detected with UPP and PetC antibodies. Thyroglobulin (669 kDa), Apoferritin from horse spleen (443 kDa) and b-Amylase from sweet potato (200 kDa) were used as marker proteins. (**D**) Maximum quantum yield of PSII of five leaves from three independent plants of Col-0 and UPP-knockdown lines (*ami-upp-4* and *ami-upp-7*) supplemented with 2.5 mM Lincomycin and illuminated with HL. (**E**) *In vivo* radiolabelling of chloroplast proteins in leave discs of Col-0 and UPP-knockdown lines (*ami-upp-4* and *ami-upp-7*) (n = 6). (**F**) Yield(II), NPQ and NO of Col-0, UPP-overexpressing line *UPP*-OX and UPP-knockdown lines (*ami-upp-4* and *ami-upp-7*) at PAR 134. Shown are measurements under control conditions floated in water (*n* = 5) compared to leave discs in 5µM DCMU for 90 min (*n* = 3). Plotted are the mean values of biological replicates +/− standard deviation. For statistical analysis two-way ANOVA (**D, E**) was performed (* = *p* < 0.05; ** = *p* < 0.01; *** = *p* < 0.001). All immunoblots and BN-PAGE were repeated at least three times, with highly similar results (**A-C**).

Next, a Blue Native-polyacrylamide gel electrophoresis was performed on thylakoids of Col-0 plants to examine a putative interaction between UPP and PetC. PetC, which is part of the cytochrome *b*_6_*f* complex, is consistently found at higher molecular weights. UPP may exist as a tetramer, with a maximum size of approximately 100 kDa. However, it was discovered that both PetC and UPP occur at the same molecular weight of around 400 kDa, indicating that they are part of the same complex (Figure 5C).

As increased levels of ribosomal proteins were observed (Supplemental Figure S3C), we quantified the translation efficiency in this organelle using two independent setups. Firstly, chloroplast translation was blocked by lincomycin. After overnight incubation in the dark, high light was applied for a further 4 hours, and the photosynthetic efficiency (Fv/Fm) was determined by pulse amplitude modulation (PAM). At all time-points, Fv/Fm was significantly reduced in *ami-upp-7*, and in *ami-upp-4* at 1 h and 4 h HL (Figure 5D). Newly synthesized proteins in leaf discs of five-week-old plants were labelled with ^35^S and their activity in total labelled proteins was quantified over 40 min. After 15 min of labelling, both *ami-upp-4* and -*7* showed significantly lower incorporation, and *ami-upp-7* also showed lower incorporation after 40 min (Figure 5E). Both experiments suggest that low UPP levels reduce chloroplast translation, which contradicts the finding of higher levels of ribosomal proteins. This observation indicates altered proteostasis, which may be caused by impaired protease activity.

These changes in the composition of Cyt *b*_6_*f* will ultimately affect photosynthetic performance, as previously described by Mainguet et al. (2009) and Ohler et al. (2019). Upon reinvestigating photosynthetic parameters in leaves, impaired yield (YII) was observed in *ami-upp-4* and -*7*, compared to Col-0 and *UPP*-OX (Figure 5F). While *ami-upp-4* compensated by increasing NPQ, *ami-upp-7* was unable to do so, resulting in unregulated energy dissipation (NO) increasing massively by 290% (Figure 5F). High NO levels similar to those induced by treatment with 3-(3,4-dichlorophenyl)-1,1-dimethylurea (DCMU), an herbicide that blocks electron transport at the Q_b_ site of PSII (Russell et al. 1995), were observed in both untreated *ami-upp-7* plants and Col-0 leaf discs treated with DCMU (Figure 5F). Additionally, a further increase in NO was observed in DCMU-treated *ami-upp-4* and -*7*. UPP-G224L was identified as a mutant not able to complement *upp-1*. When the recombinant, purified protein was subjected to size exclusion chromatography, most of the protein eluted at a size corresponding to an UPP dimer, whereas wild-type UPP appeared as tetramer (Supplemental Figure S5).

### Flavonoid biosynthesis is impaired in high light treated *ami-upp-4* line

During the testing of *UPP* mutants in various growth regimes, it was observed that *ami-upp-4* and -*7* lacked the typical red leaf coloration after several days of HL treatment. The quantification of anthocyanins revealed a 14-fold increase in Col-0 after 4d HL, which was similar in *UPP*-OX, but only a 7-fold increase in *ami-upp-4* (Figure 6A). The quantification of gene expression for early and late flavonoid biosynthetic genes, as well as the corresponding transcription factors (TFs, Araguirang and Richter 2022), revealed a significant reduction in the induction of transcripts essential for anthocyanin biosynthesis in HL-treated *ami-upp-1* and *-4* mutants (Figure 6B). The degree of transcript reduction correlated well with the reduced UPP expression and growth in the corresponding mutants. Figure 6B shows that the expression of PAP1, a major transcription factor essential for the induction of anthocyanin biosynthetic genes in HL, was reduced. This suggests downregulation of sugar-related signaling pathways (Zirngibl et al., 2022). Quantification of neutral sugars glucose, fructose, and sucrose revealed significant reductions in HL-treated *ami-upp-7*, but not in *upp-ami-4* (Figure 6C). However, both mutants were unable to increase starch levels after HL treatment compared to Col-0 (Figure 6C). Glucose-6-phosphate and fructose-6-phosphate levels were similar to those of Col-0. HL treated *ami-upp-4* showed a reduction in phosphoenolpyruvate (PEP), which is a precursor of the chloroplast shikimate pathway, as well as fumaric acid and raffinose, both of which are relevant for stress responses (Kaplan et al., 2007; Dyson et al., 2016) (Figure 6D). In the context of altered carbohydrate levels, a significant reduction in TPT levels was observed in *ami-upp-4* after 3, 8, and 24 hours of HL exposure (Figure 6E). TPT is the primary export route for photoassimilates and the most prominent membrane protein in the envelope membrane (Flügge, 1999).

**Figure 6.**
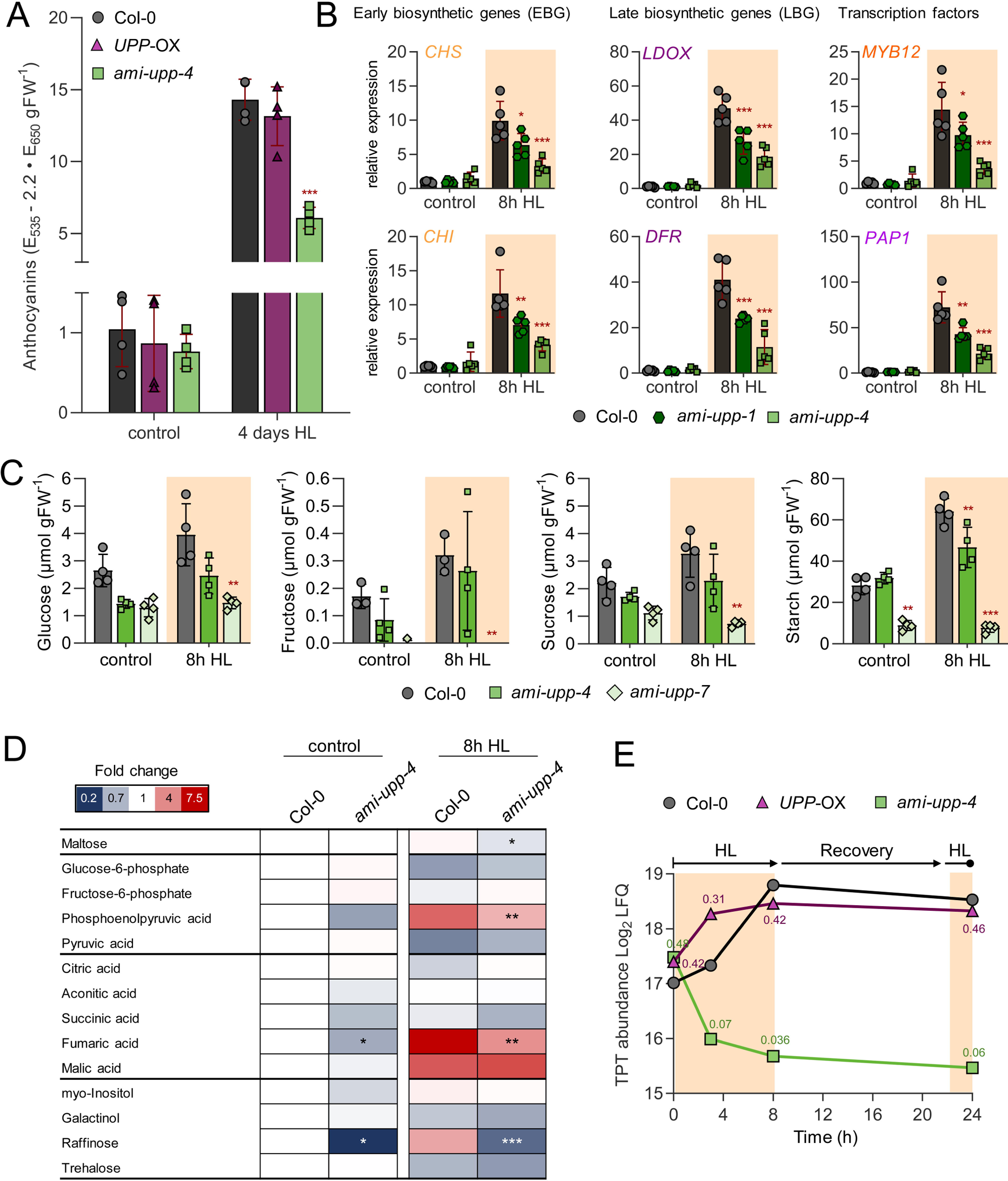
Flavonoid biosynthesis is impaired in *ami-upp* mutants. (**A**) Anthocyanin contents from five-week-old plants quantified after 4 days in high light (HL) compared to control conditions (*n* = 4). (**B**) Quantification of early (EBGs) and late biosynthetic genes (LBGs) and transcription factors of flavonoid biosynthesis Col-0, *ami-upp-1* and *ami-upp-4* after 0h and 8h of HL (1000 μmol photons m^−2^ s^−1^), determined by qRT-PCR (n = 5) relative to Col-0 (normalized to actin) (**C**) Glucose, fructose, sucrose and starch contents of Col-0, *ami-upp-4* and *ami-upp-7* after 0 and 8 hours of HL (n = 4). (**D**) Heat map of metabolite analysis of Col-0 and *ami-upp-4* after 0h and 8h HL, illustrated as fold change of *ami-upp-4* relative to Col-0 under control conditions (n = 5). (**E**) HL time course of TPT amounts from mass spec quantification (Log_2_ LFQ) of TPT (Triosephosphate phosphate translocator) from a HL time course experiment. P-values of a pairwise comparison to the corresponding Col-0 (t-test) are shown. Plotted are the mean values of biological replicates with the corresponding standard deviation. For statistical analysis one-way ANOVA (**A**), two-way ANOVA (**B**, **C**) and Students *t*-test (**D**) was performed (* = p < 0.05; ** = p < 0.01; *** = p < 0.001).

Amino acids act as the fundamental constituents of proteins, metabolic precursors, and signaling molecules. Their biosynthesis and degradation undergo changes during plant development and respond to stress levels. Typically, amino acid levels rise under stress conditions (Hausler et al., 2014) including the most abundant amino acids, glutamate, glutamine, and aspartate. The study measured amino acids in *ami-upp-4* and *UPP*-OX mutants under different conditions. It was observed that *UPP*-OX mutants showed little or no changes in amino acid levels compared to Col-0, while *ami-upp-4* mutants showed an increase in seven amino acids after 3 h HL, which became more pronounced after 8 h HL. The highest increases were measured in serine, aspartate, and glutamate (Supplemental Figure S6). During the recovery phase after the night, most amino acids in *ami-upp-4* decreased again. The largest increase was observed for alanine after 3h HL (Supplemental Figure S6).

## Discussion

### UPP affects chloroplast proteostasis and thereby impairs accumulation of PetC

The findings of this study indicate that UPP plays a crucial role in maintaining chloroplast proteostasis. Proteomics and immunoblot analysis revealed that PetC, a subunit of the cytochrome *b*_6_*f* (Cyt *b*_6_*f*) complex, is a major target. The Cyt *b*_6_*f* complex acts as a central node between the two photosystems in the thylakoid membrane of chloroplasts. Notably, the absence of PetC in *petc-2* mutants only allows growth in the presence of sucrose (Maiwald et al., 2003). Reduced PetC levels may account for the failure of *ami-upp*-lines to establish NPQ. Additionally, PetC was previously identified as an interaction partner of UPP in yeast two-hybrid studies (Arabidopsis Interactome Mapping, 2011). Blue native page followed by immunoblotting revealed that UPP and PetC run at the same molecular weight, which is significantly higher than that of tetrameric UPP. Therefore, it is likely that both proteins interact in the same complex.

Since the PetC transcript remained unchanged in *ami-upp-4* compared to Col-0, the observed decrease must take place at the protein level. This could be due to reduced translation, import into chloroplasts and thylakoids, altered turnover, or assembly into the Cyt *b*_6_*f* complex. A correlation between UPP and PetC was found in de-etiolation experiments. Here we could demonstrate that PetC levels increase significantly after 24 hours of light in Col-0, but not in *ami-upp-7*. Both proteins were present in low levels in dark-grown seedlings and increased significantly shortly after the start of illumination in our work in line with observations found in Pipitone et al. (2021). Additionally, a strong correlation was also found between UPP protein levels, PetC levels, and the growth performance of corresponding mature plants (*ami-upp* and *UPP*-OX). We interpret this as a hint for a direct role of UPP in regulating photosynthesis at the level of PetC which then affects assimilation and growth. In omics experiments, no apparent link to other metabolic or regulatory pathways that might be affected by UPP was seen. A putative obvious link to nucleotide metabolism is also less likely as nucleotide amounts were not changed in *ami-upp.* The genetic background also did not influence the increased amounts of ADP and dADP accompanied by decreased UDP levels that we observed upon HL treatment. The previously described significant increase of ATP in the cold (Chen et al., 2022) was also not altered in any of the genotypes.

UPP mutants display noticeable changes in photosynthetic energy dissipation. While *ami-upp-4* can compensate for reductions in yield (YII) by increasing NPQ, the strongest line (*ami-upp-7*) fails to do so and instead increases non-regulated energy dissipation (NO). NO increases when electron transport is blocked between PSII and the Cytb6f complex, as is the case in plants treated with 3-(3’,4’-Dichlorophenyl)-1’,1’-dimethyl urea (DCMU) (Russell et al., 1995; Vogel et al., 2012). Thus, the NO levels in *ami-upp-7* grown under control conditions are like those of DCMU-treated Col-0 plants (see Figure 6). This observation provides additional experimental evidence of a block in electron flow in cytb6/f, which regulates NPQ based on the pmf across the thylakoid (Armbruster et al., 2017) (Figure 6B).

Therefore, we conclude that UPP plays a role in stabilising or protecting PetC against degradation or supporting its repair. Little is known about this process. However, recently, de-etiolation induced protein 1 (DEIP1) was identified as an interactor of PetA and PetB subunits of Cyt *b*_6_*f*. The Cyt *b*_6_*f* core subunits PetA, B and C were missing in *deip-1* loss of function mutants (Sandoval-Ibanez et al., 2022). As previously observed for *petc-2*, *deip-1* exhibits similarities with *upp-1* and *upp-ami* lines in terms of phenotype, such as smaller habitus with pale leaves and reduced chloroplast size.

Protein amount alterations were detected in signal processing peptidases, ClpP6 protease complex subunits, and thylakoid FtsH proteases in *ami-upp-4* (Figure 4C), all of which are crucial for coordinating the maturation and degradation of the chloroplast proteome (Rowland et al., 2019). In addition, the CPN60 chloroplast chaperone complex is an essential component of the chloroplast protein folding machinery (Ries et al., 2023) and might be involved. How do proteases and chaperones interact with Cyt *b*_6_*f*? It was proposed that PetC assembles with Cpn60 in complexes before being incorporated into the Cyt *b*_6_*f* complex (Molik et al., 2001). However, this assumption was recently questioned by screening for CPN60 interactors through competition co-immunoprecipitation, where neither PetC nor UPP were identified (Ries et al., 2023). Nevertheless, interactions between the co-chaperonin lid and the ClpP protease complex were identified. The amount of ClpP subunits was found to be altered in *ami-upp-4*, and UPP was identified as a substrate of ClpP6 (Sjogren et al., 2006), suggesting the possibility of indirect interaction between these complexes and UPP.

UPP may play a role in stabilising PetC during its import into the thylakoid. PetC is imported into the thylakoids via the cpTat or delta pH pathway to be assembled into the cytochrome *b*_6_*f* complex. However, its import is slower compared to other thylakoid proteins, likely due to the need to incorporate Fe-S cofactor. UPP may bind to PetC during this process in a holdase-like manner. Holdases are typically small, abundant proteins, like the many small heat shock factors (Reinle et al., 2016) and UPP is similar in this respect. It has been shown that only oligomers, but not dimers, can exhibit holdase function for Get3 (Ulrich et al., 2023). Similarly, the UPP-G224L mutant appears as a dimer in contrast to the wildtype tetramer (Supplemental Figure S5). Although wild-type UPP and other point-mutated versions of UPP were able to complement *upp-1* (Ohler et al., 2019), we were unable to achieve complementation with UPP-G224L. This suggests that the oligomeric state of UPP as a tetramer is crucial for its function.

### UPP mutants fail to accumulate sugars and anthocyanin in HL

An interesting finding was that *ami-upp-4* and -*7* were unable to accumulate anthocyanins after HL-treatment. Upon analysing the expression of genes involved in flavonoid biosynthesis, it was observed that all tested biosynthesis genes were less induced by HL in the mutants compared to Col-0, including the transcription factors Myb12 and Pap1 (Figure 6). This implies that *ami-upp* mutants failed to induce flavonoid biosynthesis. It was demonstrated that upon HL sugars, particularly triosephosphates (TP), exported to the cytosol by TPT, are crucial for this process (Zirngibl et al., 2022). In line with this, the detectable amount of TPT protein in *ami-upp-4* was significantly reduced already after 3h HL and even more so after 8h HL. Therefore, it is probable that the reduced export of TP from the chloroplast and the absence of the inducing sugar signal are responsible for the decreased anthocyanin biosynthesis in UPP mutants.

The photosynthetic efficiency of *ami-upp-4* is reduced, but this is largely compensated by the reduced growth. Neutral sugars and hexose phosphates were not significantly altered in these plants. However, starch accumulation in HL was significantly reduced (Figure 6). Reduced TPT amounts may function to retract triosephosphates in the chloroplast to prevent depletion of the CBB or to support the oxidative pentose phosphate pathway in the dark. Protease activity is most likely required to achieve such a fast a reduction of more than 8-fold in TPT, the most abundant protein in the chloroplast envelope (Flügge, 1999). This phenomenon has not been previously described. Future studies should investigate whether the rhomboid-like proteases RBL10 and RBL11. So far, the only known proteases acting in the chloroplast envelope (Lavell et al., 2019) are involved in regulating TPT abundance.

Besides anthocyanins, raffinose, phosphoenolpyruvate and fumarate were significantly reduced in *ami-upp-4* under control and HL conditions compared to Col-0 (Figure 6). Increased levels of these metabolites are a typical HL response and this is also reflected in our analysis in Col-0 plants. Fumarate and starch can act as flexible and alternative carbon sinks in Arabidopsis and FUM2 is a key regulator in this scenario (Chia et al., 2000; Pracharoenwattana et al., 2010). In line with reduced levels of fumarate, we observed decreased levels of FUM2, which may be attributed to the stabilization of starch levels in *ami-upp-4*. Moreover, the inability of *ami-upp-4* to accumulate these osmoprotectants is most likely a consequence of reduced photosynthetic capacity in these mutants.

## Materials and Methods

### Plant growth

Plant growth was performed according to the protocol described in Ohler et al. (2019), with the following modifications: normal light intensity was set to 120 μmol photons m^−2^ s^−1^, high light intensity was set to 1000 μmol photons m^−2^ s^−1^ at 21°C. Cold experiments were performed at 4°C for 24 hours at normal light intensity. Samples for metabolomics (six independent biological replicates) and proteomics (three independent biological replicates obtained by combining two samples from the metabolomic analysis) were collected from five-week-old plants. The rosette material was harvested and immediately frozen in liquid nitrogen. De-etiolation experiments were conducted following the procedure outlined in Pipitone et al. (2023). After imbibition and growth for seven days in complete darkness, the seedlings were grown under constant light (40 μmol photons m^−2^ s^−1^) for 24 hours.

### Generation of *UPP* knock-down plants

The protocol of Schwab et al. (2006) was used to generate knock-down mutants of UPP (At3g53900) through gene silencing with artificial microRNA (*amiRNA*). Different primers with Gateway™-compatible sequences attP1 and attP2 (Supplemental Table S2) were designed using an online tool (www.weigelworld.com) to target UPP. The cloned fragments were subcloned into the Gateway™ entry vector pDONR/Zeo via a BP clonase reaction. Later, they were subcloned into the target vector pK2GW7 downstream of a 35S-CaMV promoter via an LR clonase reaction. Transformation of Arabidopsis thaliana Wildtype (Col-0) plants was carried out by floral dip (Narusaka et al., 2010) using previously transformed Agrobacterium tumefaciens GV3101 (Furini et al., 1994). Positive transformed plants were initially selected on ½ MS agar plates supplemented with Kanamycin and subsequently transferred to soil. The UPP transcript levels were quantified to identify different independent lines with 54 - 4.6% UPP transcript remaining.

### Transcript quantification with qRT PCR

RNA isolation, cDNA preparation, and PCR were performed following the protocol described in Ohler et al. (2019). Briefly, RNA was isolated from whole plant tissue ground in liquid nitrogen using the Nucleospin RNA Plant Kit (Macherey-Nagel, Düren, Germany) according to the manufacturer’s instructions. The qScript cDNA Synthesis Kit (Quantabio, United States) was used for RNA transcription into cDNA. Transcripts related to flavonoid biosynthesis and corresponding primers for qRT-PCR are described in (Zirngibl et al., 2022). Additional primers for UPP are listed in Supplemental Table S2.

### Protein extraction, gel-electrophoresis and immunoblotting

Proteins were extracted according to (Ermakova et al., 2019). Therefore, 50-100 mg leaf samples of were grinded in liquid nitrogen with protein extraction buffer (100mM trisaminomethane-HCl, pH 7.8, supplemented with 25 mM NaCl, 20mM ethylenediaminetetraacetic acid, 2% sodium dodecyl sulfate (w/v), 10 mM dithiothreitol and 2% (v/v) protease inhibitor cocktail (Sigma, St Louis, MO). Protein extracts were incubated at 65 °C for 10 min and then centrifuged at 13,000 g for 1 min at 4 °C. Protein extracts were supplemented with 4× Laemmli buffer (BioRad, Hercules, CA), the samples were stored at −80°C until useSDS page was performed with BioRad gels and according to the manufacturers protocol.

Immunoblots were essentially performed as given in (Ohler et al., 2019), here the proteins were separated on SDS-gels and transferred onto a nitrocellulose membrane by the semi-dry Transblot Turbo Transfer System (Bio-Rad, Hercules, CA, USA). Primary antibodies were obtained from Agrisera (Vännäs, Sweden) except for anti-UPP (Ohler et al., 2019). For blue native electrophoresis (BN-PAGE), extraction of thylakoids and was done as given in (Schägger et al., 1994). After electrophoresis, the gels were blotted onto a PVDF membrane as described above.

### Photosynthetic activity was measured using pulse-amplitude-modulation

(PAM) fluorometry, as previously described (Bellin et al., 2023). Five-week-old plants grown under standard conditions were transferred to high light, and leaf discs from respective plants were treated with DCMU (5µM). Five leaf discs were incubated for 90 minutes in 5ml buffer with CaCl_2_ (Russell et al., 1995).

### Statistical analyses

Statistical analysis of qRT-PCR, proteomic and metabolomic analysis data was performed using the Student’s *t*-test (see also legend for details). Asterisks indicate statistically significant differences determined using Student’s *t*-test (\**p* < 0.05; \*\**p* < 0.01, \*\*\**p* < 0.001). The significance criteria for comparisons within the proteomic dataset were based on adjusted *p* < 0.05. Changes in protein levels within a given pairwise comparison were expressed as log2 fold change (log 2 FC), or as fold change (FC) where FC equals the second condition divided by the first condition (Pagano et al., 2023).

### 35S pulse labelling of proteins in leaves

Labelling of proteins in leaf discs by ^35^S methionine/cysteine was performed as given in (DeTar et al., 2021).

## Acknowledgement and funding

This work was funded by DFG grants (CRC Transregio TRR175, B06 to S.S. B08 to T.M, C06 to A.R. and C05 to D.L.).

## Author contributions

T.M. conceived and supervised the study, obtained funding, and provided resources. V.S. generated all mutants and performed characterization, and phenotyping. V.S., L.B. performed analysis and interpretation of proteomic and metabolomic data. S.S. performed proteomics. M.L. performed metabolomic data analysis. D.L. supervised proteomics and metabolomics J.R. performed and C.P.W. advised and interpreted nucleotide quantification. K.J. performed and A.R. advised and interpreted expression of flavonoid biosynthesis genes. T.P. supported immunoblotting and quantification A.L. supported results interpretation related to proteostasis, L.O. and S.S. performed UPP mutagenesis, SEC and complementation trials. T.M., V.S. and L.B. wrote the original draft. All authors reviewed and agreed to the final manuscript.

## Data Availability statement

Transcriptome data from next generation sequencing were deposited in the European Nucleotide Archive under Accession number: (xxxxxxx).

Proteomic data are deposited in the PRIDE Archive under Accession number: (xxxxxx)

## References

Arabidopsis Interactome Mapping Consortium (2011) Evidence for network evolution in an Arabidopsis interactome map. Science 333: 601–607

Araguirang GE, Richter AS (2022) Activation of anthocyanin biosynthesis in high light - what is the initial signal? New Phytol 236: 2037–2043

Armbruster U, Correa Galvis V, Kunz HH, Strand DD (2017) The regulation of the chloroplast proton motive force plays a key role for photosynthesis in fluctuating light. Curr Opin Plant Biol 37: 56–62

Arrivault S (2019) UMP Pyrophosphorylase: A moonlighting protein with essential functions in chloroplast development and photosynthesis establishment. Plant Physiol 180: 1779–1780

Bae W, Lee YJ, Kim DH, Lee J, Kim S, Sohn EJ, Hwang I (2008) AKR2A-mediated import of chloroplast outer membrane proteins is essential for chloroplast biogenesis. Nat Cell Biol 10: 220–227

Bellin L, Del Cano-Ochoa F, Velazquez-Campoy A, Möhlmann T, Ramón-Maiques S (2021) Mechanisms of feedback inhibition and sequential firing of active sites in plant aspartate transcarbamoylase. Nat Commun 12: 947

Bellin L, Garza Amaya DL, Scherer V, Pruss T, John A, Richter A, Möhlmann T (2023) Nucleotide imbalance, provoked by downregulation of Aspartate Transcarbamoylase impairs cold acclimation in Arabidopsis. Molecules 28

Bellin L, Scherer V, Dörfer E, Lau A, Vicente AM, Meurer J, Hickl D, Möhlmann T (2021) Cytosolic CTP production limits the establishment of photosynthesis in Arabidopsis. Front Plant Sci 12: 789189

Chen X, Kim SH, Rhee S, Witte CP (2023) A plastid nucleoside kinase is involved in inosine salvage and control of purine nucleotide biosynthesis. Plant Cell 35: 510–528

Chia DW, Yoder TJ, Reiter WD, Gibson SI (2000) Fumaric acid: an overlooked form of fixed carbon in Arabidopsis and other plant species. Planta 211: 743–751

Clausen AR, Girandon L, Ali A, Knecht W, Rozpedowska E, Sandrini MP, Andreasson E, Munch-Petersen B, Piskur J (2012) Two thymidine kinases and one multisubstrate deoxyribonucleoside kinase salvage DNA precursors in Arabidopsis thaliana. FEBS J 279: 3889–3897

Correa-Galvis V, Poschmann G, Melzer M, Stuhler K, Jahns P (2016) PsbS interactions involved in the activation of energy dissipation in Arabidopsis. Nat Plants 2: 15225

DeTar RA, Barahimipour R, Manavski N, Schwenkert S, Hohner R, Bölter B, Inaba T, Meurer J, Zoschke R, Kunz HH (2021) Loss of inner-envelope K^+^/H^+^ exchangers impairs plastid rRNA maturation and gene expression. Plant Cell 33: 2479–2505

Dorion S, Rivoal J (2015) Clues to the functions of plant NDPK isoforms. Naunyn-Schmiedeberg’s archives of pharmacology 388: 119–132

Dyson BC, Miller MA, Feil R, Rattray N, Bowsher CG, Goodacre R, Lunn JE, Johnson GN (2016) FUM2, a cytosolic fumarase, is essential for acclimation to low temperature in Arabidopsis thaliana. Plant Physiol 172: 118–127

Ermakova M, Lopez-Calcagno PE, Raines CA, Furbank RT, von Caemmerer S (2019) Overexpression of the Rieske FeS protein of the Cytochrome b(6)f complex increases C(4) photosynthesis in *Setaria viridis*. Commun Biol 2: 314

Espinoza-Corral R, Schwenkert S, Schneider A (2023) Characterization of the preferred cation cofactors of chloroplast protein kinases in Arabidopsis thaliana. FEBS Open Bio 13: 511–518

Flügge UI (1999) Phosphate translocators in plastids. Annu Rev Plant Physiol Plant Mol Biol 50: 27–45

Furini A, Koncz C, Salamini F, Bartels D (1994) Agrobacterium-mediated transformation of the desiccation-tolerant plant *Craterostigma-Plantagineum*. Plant Cell Reports 14: 102–106

Häusler RE, Ludewig F, Krueger S (2014) Amino acids--a life between metabolism and signaling. Plant Sci 229: 225–237

Jarvi S, Suorsa M, Tadini L, Ivanauskaite A, Rantala S, Allahverdiyeva Y, Leister D, Aro EM (2016) Thylakoid-Bound FtsH Proteins Facilitate Proper Biosynthesis of Photosystem I. Plant Physiol 171: 1333–1343

Kato Y, Hyodo K, Sakamoto W (2018) The Photosystem II Repair Cycle Requires FtsH Turnover through the EngA GTPase. Plant Physiol 178: 596–611

Kaplan F, Kopka J, Sung DY, Zhao W, Popp M, Porat R, Guy CL (2007) Transcript and metabolite profiling during cold acclimation of Arabidopsis reveals an intricate relationship of cold-regulated gene expression with modifications in metabolite content. Plant J 50: 967–981

Lavell A, Froehlich JE, Baylis O, Rotondo AD, Benning C (2019) A predicted plastid rhomboid protease affects phosphatidic acid metabolism in *Arabidopsis thaliana*. Plant J 99: 978–987

Li L, Wurtele ES (2015) The QQS orphan gene of Arabidopsis modulates carbon and nitrogen allocation in soybean. Plant Biotechnol J 13: 177–187

Mainguet SE, Gakiere B, Majira A, Pelletier S, Bringel F, Guerard F, Caboche M, Berthome R, Renou JP (2009) Uracil salvage is necessary for early Arabidopsis development. Plant J 60: 280–291

Maiwald D, Dietzmann A, Jahns P, Pesaresi P, Joliot P, Joliot A, Levin JZ, Salamini F, Leister D (2003) Knock-out of the genes coding for the Rieske protein and the ATP-synthase delta-subunit of Arabidopsis. Effects on photosynthesis, thylakoid protein composition, and nuclear chloroplast gene expression. Plant Physiol 133: 191–202

Malnoe A, Wang F, Girard-Bascou J, Wollman FA, de Vitry C (2014) Thylakoid FtsH protease contributes to photosystem II and cytochrome b6f remodeling in *Chlamydomonas reinhardtii* under stress conditions. Plant Cell 26: 373–390

Molik S, Karnauchov I, Weidlich C, Herrmann RG, Klösgen RB (2001) The Rieske Fe/S protein of the cytochrome b6/f complex in chloroplasts: missing link in the evolution of protein transport pathways in chloroplasts? J Biol Chem 276: 42761–42766

Narusaka M, Shiraishi T, Iwabuchi M, Narusaka Y (2010) The floral inoculating protocol: a simplified *Arabidopsis thaliana* transformation method modified from floral dipping. Plant Biotechnology 27: 349–351

Ohler L, Niopek-Witz S, Mainguet SE, Möhlmann T (2019) Pyrimidine salvage: physiological functions and interaction with chloroplast biogenesis. Plant Physiol 180: 1816–1828

Pagano A, Kunz L, Dittmann A, Araujo SS, Macovei A, Shridhar Gaonkar S, Sincinelli F, Wazeer H, Balestrazzi A (2023) Changes in *Medicago truncatula* seed proteome along the rehydration-dehydration cycle highlight new players in the genotoxic stress response. Front Plant Sci 14: 1188546

Pipitone R, Eicke S, Pfister B, Glauser G, Falconet D, Uwizeye C, Pralon T, Zeeman SC, Kessler F, Demarsy E (2021) A multifaceted analysis reveals two distinct phases of chloroplast biogenesis during de-etiolation in Arabidopsis. Elife 10

Pracharoenwattana I, Zhou W, Keech O, Francisco PB, Udomchalothorn T, Tschoep H, Stitt M, Gibon Y, Smith SM (2010) Arabidopsis has a cytosolic fumarase required for the massive allocation of photosynthate into fumaric acid and for rapid plant growth on high nitrogen. Plant J 62: 785–795

Ries F, Weil HL, Herkt C, Mühlhaus T, Sommer F, Schroda M, Willmund F (2023) Competition co-immunoprecipitation reveals the interactors of the chloroplast CPN60 chaperonin machinery. Plant Cell Environ 46: 3371–3391

Russell AW, Critchley C, Robinson SA, Franklin LA, Seaton G, Chow WS, Anderson JM, Osmond CB (1995) Photosystem II regulation and dynamics of the chloroplast D1 protein in Arabidopsis leaves during photosynthesis and photoinhibition. Plant Physiol 107: 943–952

Schägger H, Cramer WA, von Jagow G (1994) Analysis of molecular masses and oligomeric states of protein complexes by blue native electrophoresis and isolation of membrane protein complexes by two-dimensional native electrophoresis. Anal Biochem 217: 220–230

Schmid LM, Ohler L, Möhlmann T, Brachmann A, Muino JM, Leister D, Meurer J, Manavski N (2019) PUMPKIN, the sole plastid UMP kinase, associates with group II introns and alters their metabolism. Plant Physiol 179: 248–264

Schwab R, Ossowski S, Riester M, Warthmann N, Weigel D (2006) Highly specific gene silencing by artificial microRNAs in Arabidopsis. Plant Cell 18: 1121–1133

Sjögren LL, Stanne TM, Zheng B, Sutinen S, Clarke AK (2006) Structural and functional insights into the chloroplast ATP-dependent Clp protease in Arabidopsis. Plant Cell 18: 2635–2649

Stitt M, Lilley R, Gerhardt R, Heldt HW (1989) Metabolite levels in specific cells and subcellular compartments of plant leaves. Meth. Enzymol 174: 518–552

Straube H, Niehaus M, Zwittian S, Witte CP, Herde M (2021) Enhanced nucleotide analysis enables the quantification of deoxynucleotides in plants and algae revealing connections between nucleoside and deoxynucleoside metabolism. Plant Cell 33: 270–289

Thimm O, Bläsing O, Gibon Y, Nagel A, Meyer S, Kruger P, Selbig J, Muller LA, Rhee SY, Stitt M (2004) MAPMAN: a user-driven tool to display genomics data sets onto diagrams of metabolic pathways and other biological processes. Plant J 37: 914–939

Tyanova S, Temu T, Sinitcyn P, Carlson A, Hein MY, Geiger T, Mann M, Cox J (2016) The Perseus computational platform for comprehensive analysis of (prote)omics data. Nat Methods 13: 731–740

Tzvetkova-Chevolleau T, Franck F, Alawady AE, Dall’Osto L, Carriere F, Bassi R, Grimm B, Nussaume L, Havaux M (2007) The light stress-induced protein ELIP2 is a regulator of chlorophyll synthesis in Arabidopsis thaliana. Plant J 50: 795–809

Vogel MO, Gomez-Perez D, Probst N, Dietz KJ (2012) Combinatorial signal integration by APETALA2/Ethylene Response Factor (ERF)-transcription factors and the involvement of AP2-2 in starvation response. Int J Mol Sci 13: 5933–5951

Zirngibl ME, Araguirang GE, Kitashova A, Jahnke K, Rolka T, Kuhn C, Nägele T, Richter AS (2022) Triose phosphate export from chloroplasts and cellular sugar content regulate anthocyanin biosynthesis during high light acclimation. Plant Commun: 100423

