## Supplemental Figures 1-6 for "UPP affects chloroplast development by interfering with chloroplast proteostasis"

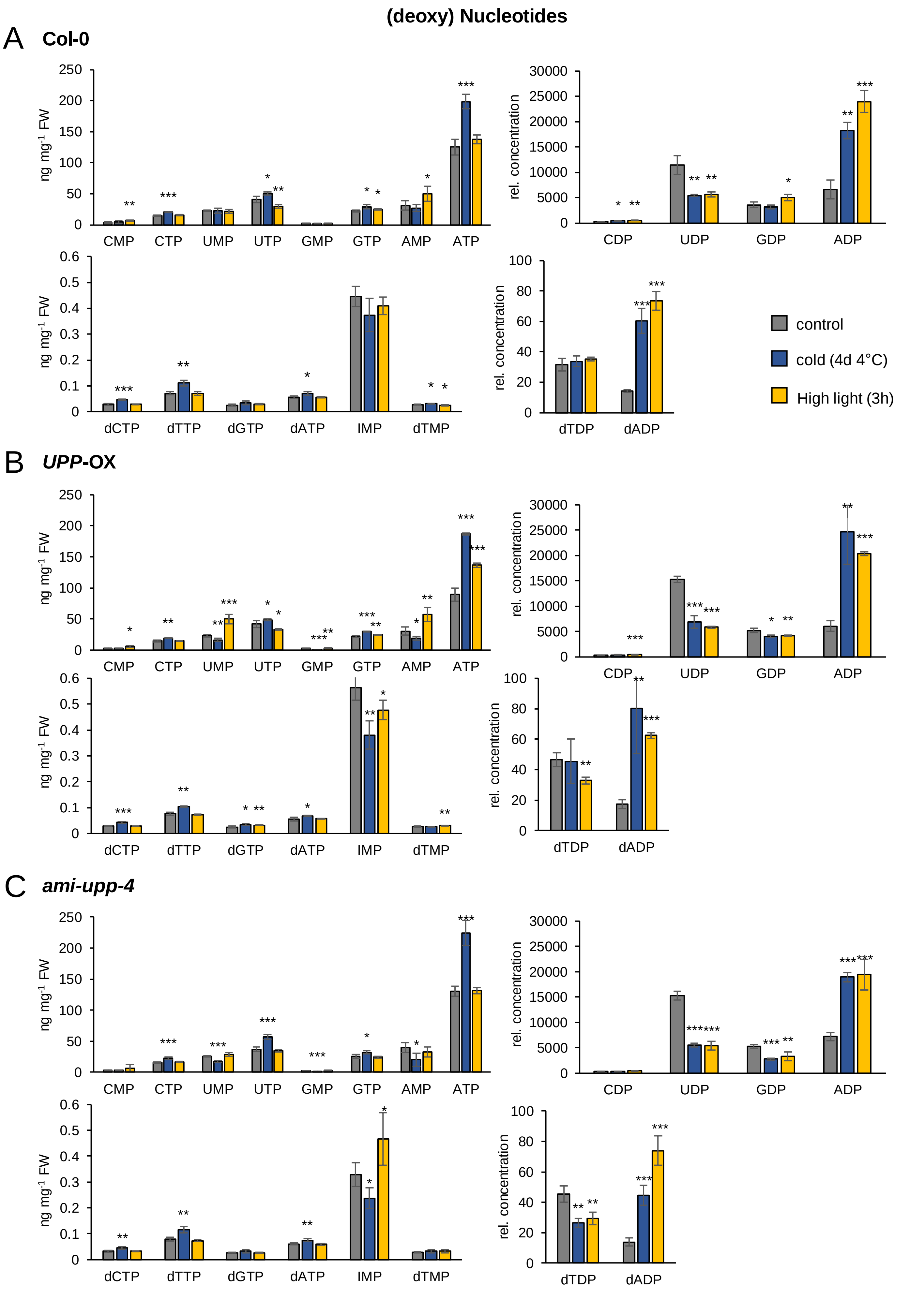


**Supplemental Figure S1. (Deoxy-) Nucleotides compared in Col-0, *UPP*-OX and *ami-upp-4* plants under control, cold and high light treatment.**

**Supplemental Figure S1. (Deoxy-) Nucleotide contents in Col-0, *UPP*-OX and *ami-upp-4* plants under control, cold and high light treatment.** (**A, B, C**) Nucleotide-mono-phosphate (NMPs), Nucleotide-di-phosphate (NDPs), nucleotide-triphosphate (NTPs) and deoxy-nucleotide contents in five-week-old (**A**) Col-0, (**B**) UPP overexpressor *UPP*-OX and (**C**) UPP knockdown *ami-upp-4* plants. Plotted are the mean values of four biological replicates with the corresponding standard deviation. For statistical analysis control treatments were compared to 4°C and high light treatments using Student's t-test (* = $p < 0.05$; ** = $p < 0.01$; *** = $p < 0.001$).

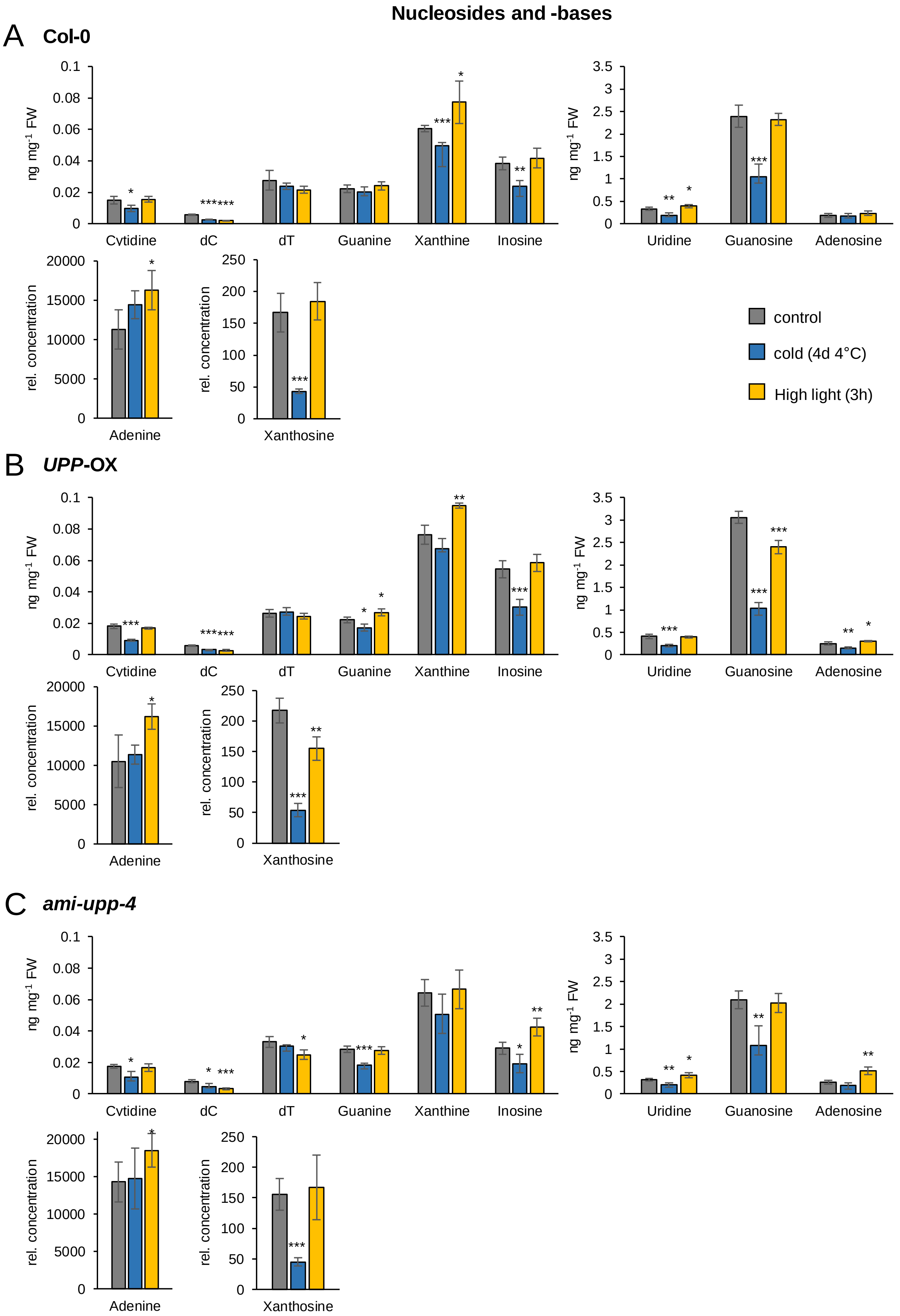


**Supplemental Figure S2. Nucleosides and Nucleobases compared in Col-0, *UPP*-OX and *ami-upp-4* plants under control, cold and high light treatment.**

**Supplemental Figure S2. Nucleoside and Nucleobase contents in Col-0, *UPP*-OX and *ami-upp-4* plants under control, cold and high light treatment.** (**A, B, C**) Nucleosides and nucleobase contents in five-week-old (**A**) Col-0, (**B**) UPP overexpressor (*UPP*-OX) and (**C**) UPP knockdown (*ami-upp-4*) plants. dC (Deoxycytidine); dT (Deoxythymidine) Plotted are the mean values of four biological replicates with the corresponding standard deviation. For statistical analysis control treatments were compared to 4°C and high light treatments using Student's t-test (* = $p < 0.05$; ** = $p < 0.01$; *** = $p < 0.001$).

A

Fold change: -0.14 | -0.07 | 0 | 0.3 | 0.57

*ami-upp-4* / Col-0

| Calvin-Benson-Bassham cycle | | 0h | 3h | 8h | Rec. |
|---|---|---|---|---|---|
| ATCG00490 | RbcL | | | * | |
| AT5G38420 | RbcS | | *** | ** | |
| AT3G12780 | PGK1 | | | * | |
| AT1G56190 | PGK1 | ** | | | |
| AT1G42970 | GapB | | | ** | |
| AT3G26650 | GapA1 | | | * | |
| AT1G12900 | GapA2 | | | * | |
| AT3G62410 | CP12 | * | | | |
| AT4G38970 | Aldolase | | | * | |
| AT4G26530 | Aldolase | | ** | ** | ** |
| AT3G54050 | FBPase | | | * | |
| AT3G55800 | SBPase | | | | * |
| AT3G60750 | TK | | | ** | |
| AT5G61410 | RPE | | | ** | * |
| AT2G28000 | CPN60A | | | | * |
| AT1G55490 | CPN60B | | * | | * |

B

Fold change: -1.5 | -0.75 | 0 | 0.15 | 0.33

*ami-upp-4* / Col-0

| Redox | | 0h | 3h | 8h | Rec. |
|---|---|---|---|---|---|
| AT3G02730 | TRXF1 | | * | * | |
| AT1G03680 | TRXM1 | | * | ** | * |
| AT5G42980 | TRX3 | | | ** | * |
| AT1G45145 | TRX5 | * | * | | * |
| AT3G54960 | PID1 | | | *** | |
| AT5G60640 | PID2 | | | ** | |
| AT1G21750 | PDI5 | | | ** | |
| AT1G77510 | PDI6 | | | ** | |
| AT2G47470 | PDIL2-1 | * | | ** | |
| AT4G03520 | ATHM | | | * | * |
| AT3G15360 | ATHM4 | | | ** | |
| AT2G41680 | NTRC | | | | ** |
| AT1G50320 | ATHX | | | * | |
| AT1G43560 | ATY2 | | * | | * |
| AT2G24940 | MAPR2 | | ** | ** | ** |
| AT3G48890 | MAPR3 | | * | | * |
| AT3G06730 | TRXfamily prot | | * | | |
| AT4G35000 | APX3 | | | * | |
| AT1G77490 | TAPX | | | * | |
| AT4G08390 | SAPX | | * | | |
| AT3G52880 | MDAR1 | | * | | |
| AT5G03630 | MDAR2 | | | * | *** |
| AT5G16710 | DHAR3 | | * | ** | |
| AT1G07890 | APX1 | | | ** | ** |
| AT1G19570 | DHAR1 | | | ** | *** |
| AT5G28840 | GME | | | ** | |
| AT1G76080 | CDSP32 | * | * | | |
| AT5G27380 | GSH2 | | | | ** |
| AT3G54660 | GR2 | | | * | |
| AT2G25080 | GPX1 | | * | | * |
| AT2G31570 | GPX2 | * | | | |
| AT4G11600 | GPX6 | | * | | |
| AT5G20140 | SOUL | | | * | |
| AT3G10520 | GLB2 | | | * | |
| AT5G20500 | ATHM | | | ** | |
| AT2G20270 | GR-put. | | | * | |
| AT5G40370 | GR-put. | | | ** | |
| AT1G65980 | TPX1 | | | ** | |
| AT5G08410 | FTRA2 | | ** | | |
| AT2G04700 | FTRx | | * | | |
| AT3G19010 | 2OG-Fe | | ** | | |
| AT5G06290 | 2cysPerxB | | * | | |
| AT3G52960 | PRX2 | | | * | |
| AT3G06050 | PRXIIF | * | | * | |
| AT4G35090 | CAT2 | | * | *** | |
| AT3G10920 | MSD1 | | | * | |
| AT1G20620 | CAT3 | * | ** | ** | *** |
| AT5G20080 | NADH-cytb5 | | * | *** | * |

C

Fold change: -1.9 | -0.9 | 0 | 0.2 | 0.44

*ami-upp-4* / Col-0

| | | | 0h | 3h | 8h | Rec. |
|---|---|---|---|---|---|---|
| Nuclear encoded ribosomes | AT1G79850 | RPS17 | | | ** | * |
| | AT2G33800 | RPS5 | | | ** | ** |
| | AT1G64510 | RPS6 | | | * | * |
| | AT1G74970 | RPS9 | | | * | |
| | AT3G15190 | RPS20 | * | | | |
| | AT5G24490 | - | | ** | | |
| | AT1G32990 | RPL11 | | | ** | ** |
| | AT1G78630 | RPL13 | * | | * | |
| | AT1G48350 | RPL18 | | | * | |
| | AT5G47190 | | * | | ** | |
| | AT1G35680 | RPL21 | | | * | ** |
| | AT5G54600 | RPL24 | | | * | * |
| | AT2G43030 | RPL3A | * | | ** | * |
| | AT1G75350 | RPL31 | ** | | ** | * |
| | AT1G07320 | RPL4 | | | ** | |
| | AT4G01310 | RPL5 | | | *** | |
| | AT1G05190 | RPL6 | | | ** | |
| | AT3G44890 | RPL9 | | | * | ** |
| | AT3G63490 | RPL1 | ** | | ** | * |
| | AT5G13510 | RPL10 | | | * | * |
| | AT3G25920 | RPL15 | | | ** | ** |
| Chloroplast encoded | ATCG00330 | RPS14 | | | | * |
| | ATCG00750 | RPS11 | | | *** | * |
| | ATCG00650 | RPS18 | | | | * |
| | ATCG00820 | RPS19 | | | ** | * |
| | ATCG00800 | RPS3 | | | ** | * |
| | ATCG00380 | RPS4 | | | | * |
| | ATCG00770 | RPS8 | | | * | |
| | ATCG00640 | RPL33 | | | * | |
| | ATCG00780 | RPL14 | | | * | ** |
| | ATCG00790 | RPL16 | | | * | * |
| | ATCG00660 | RPL20 | | | * | |

**Supplemental Figure S3. Proteomics indicate alterations in proteins involved in functional categories.** Comparison of protein alterations between Col-0 and *ami-upp-4* associated to Calvin-Benson-Bassham cycle (**A**), chloroplast ribosomes (**B**) and redox reactions (**C**) (p-value < 0.05). For statistical analysis *ami-upp-4* proteins were compared to Col-0 proteins pairwise in the same condition using Students t-test (**A**, **B**, **C**) (* = p < 0.05; ** = p < 0.01; *** = p < 0.001).

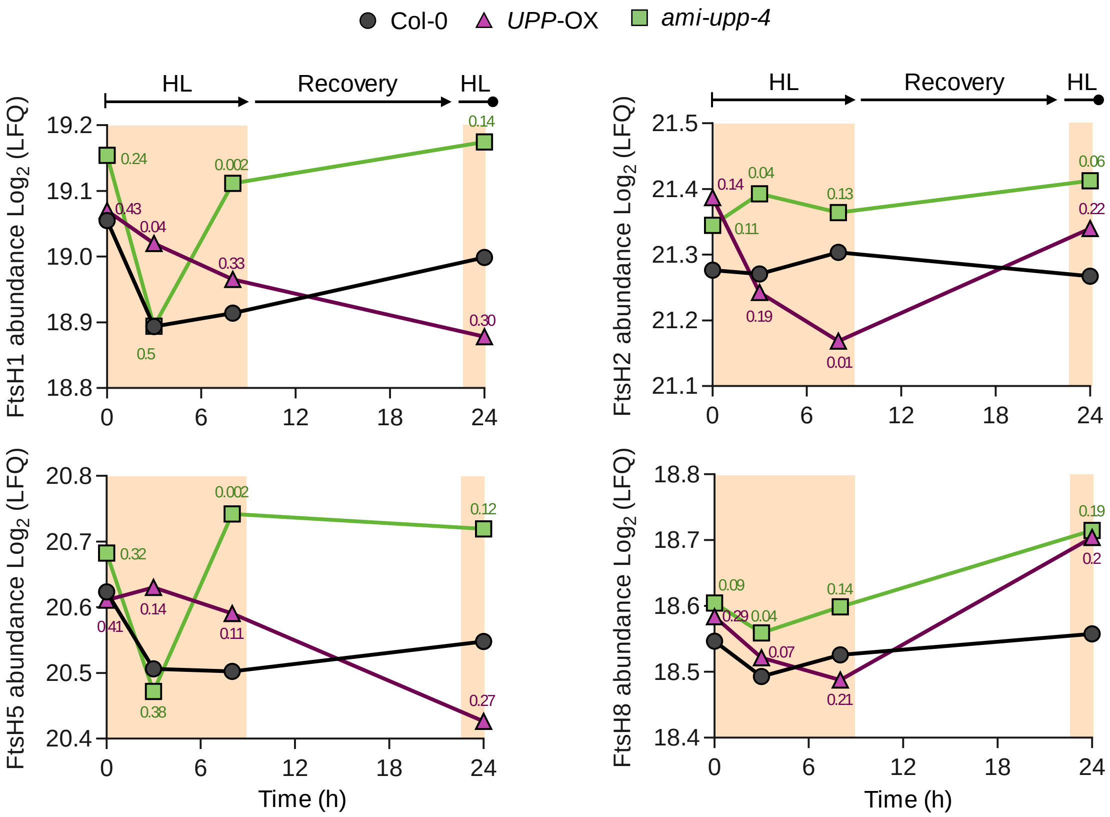


**Supplemental Figure S4. Thylakoid proteases (FtsH) are upregulated in *ami-upp* mutants.** Proteomic data are shown as $Log_2$ LFQ of FtsH proteases 1, 2, 5, and 8 of Col-0, UPP- overexpressing Line (*UPP*-OX) and UPP-knockdown Line (*ami-upp-4*). Plotted are the mean values of three biological replicates. Pairwise comparisons (*ami-upp-4* to Col-0) or (*UPP*-OX to Col-0) were analyzed by Students t-test, and p-values given for each comparison group next to the genotype symbol.

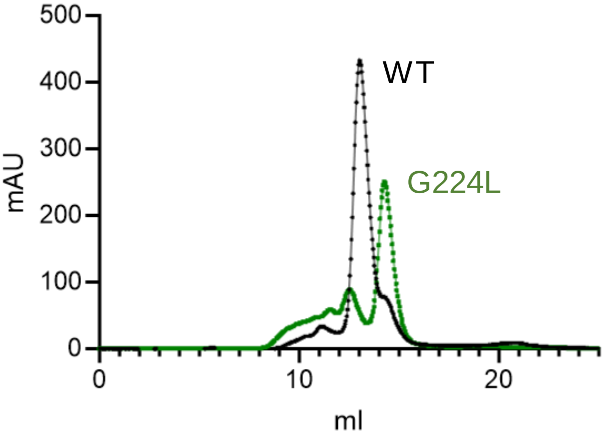


**Supplemental Figure S5. Typical size exclusion chromatogram of WT-UPP and mutated UPP (G224L) is shown** Recombinant proteins were purified from *E. coli* extracts by i-MAC as given in Ohler et al. (2019). Calibration to dextran blue and marker proteins (alcohol dehydrogenase, BSA and carboanhydrase) revealed a molecular mass of 125 kDa for UPP-WT and 69 kDa for UPP-G224L. The analysis was repeated with the same result.

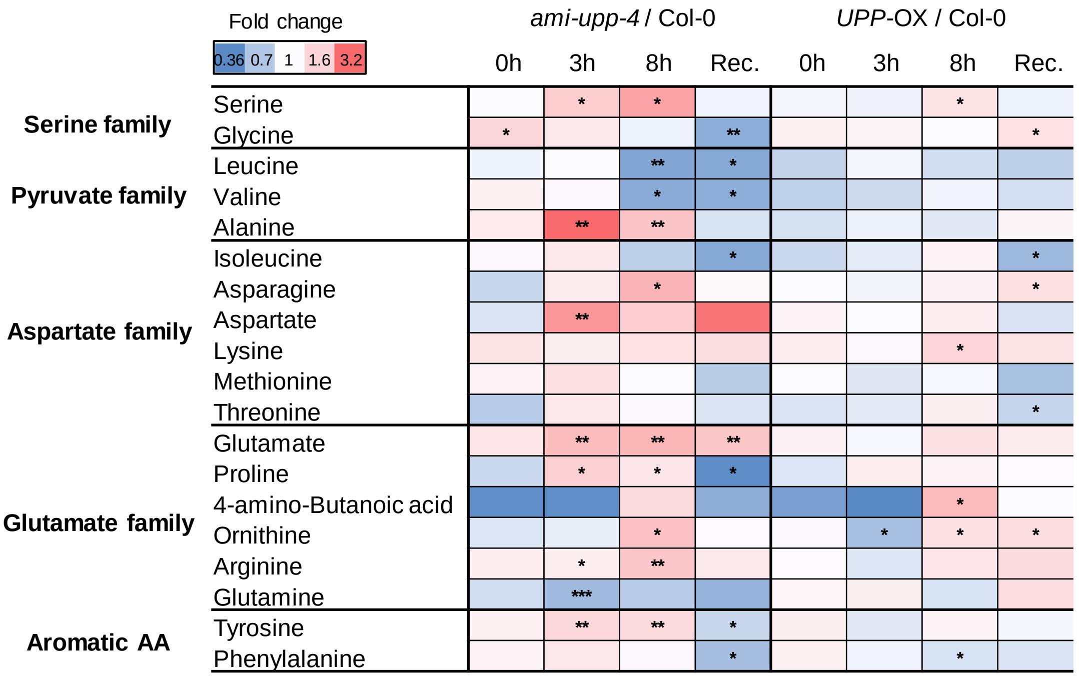


**Supplemental Figure S6. Knock-down of UPP leads to altered levels of free amino acids under HL.** Heat map of the metabolite analysis of Col-0, *UPP*-OX and *ami-upp-4* after 0h, 3h and 8h HL and recovery, illustrated as fold change of *ami-upp-4* relative to Col-0 and *UPP*-OX relative to Col-0 under the same conditions. For statistical analysis students t-test was performed (* = $p < 0.05$; ** = $p < 0.01$; *** = $p < 0.001$) (n=5 biological replicates).
