## Supplemental Methods for "UPP affects chloroplast development by interfering with chloroplast proteostasis"

**Supplemental material and Methods**

**Quantification of, anthocyanins and metabolites**

Plant material from five-week-old soil-grown plants was used. Anthocyanins were extracted from leaf tissue ground in liquid nitrogen. 50 mg of ground material was extracted in 1 ml of extraction buffer (H2O: 1-propanol: HCl (81:18:1)). Samples of four biological replicates were boiled at 95°C for 3 minutes, incubated in the dark overnight at room temperature, and the supernatant was measured photometrically at 535 nm and 650 nm. The extinction was corrected using Rayleigh's formula (E535-2.2E650).

To extract sugars and starch, 50 mg of leaf tissue was ground in liquid nitrogen. The material was then boiled in 500 µL of deionized water for 15 minutes at 95°C. The resulting supernatant was used for sugar measurements, while the pellet was used for starch measurements. Quantification was performed photometrically using a NAD+-coupled enzymatic assay (Stitt et al., 1989).

Nucleotide quantification was performed as described in (Straube et al., 2021).

The analysis of primary metabolites was carried out using GC-TOF-MS, as previously described in (Garcia-Molina et al., 2020). To do this, 100 mg of fresh plant material was ground and mixed with 180 µl of methanol at -20°C. Internal standards, ribitol (10 µl at 0.2 mg $ml^{-1}$ in water) and 13C-sorbitol (10 µl at 0.2 mg $ml^{-1}$ in water), were added to enable relative quantification of metabolites. After incubating the mixture at 70°C for 15 minutes, combine it with 100 µl of chloroform and 200 µl of water. To separate polar and non-polar compounds, undergo a 15-minute centrifugation at 25000 g for phase separation. Resuspend the resulting pellet of 50 µl of the upper (polar) phase in 10 µl of methoxyaminhydrochloride (20 mg ml-1 in pyridine). Then, 20 µl of BSTFA (N,O-Bis[trimethylsilyl]trifluoroacetamide) containing 2.5 µl of a retention time standard mixture of linear alkanes for retention time alignment was added. The mixture was shaken at 40°C for 90 minutes. The sample was then incubated at 40°C for 45 minutes after adding 20 µl of BSTFA (N,O-Bis[trimethylsilyl]trifluoroacetamide) containing 2.5 µl of a retention time standard mixture of linear alkanes for retention time alignment. GC-TOF-MS analysis was performed using a Pegasus HT system (Leco, St Joseph, USA) with an autosampler system (Combi PAL, CTC Analytics AG, Zwingen, Switzerland) after injecting 1 µl of each derivatized sample. Helium gas was used as the carrier and maintained at a constant flow rate of 1 ml/min. The system employed an Agilent 7890A oven (Agilent, Santa Clara, USA) with a VF-5ms column (30 m) and an EZ-Guard column (10 m). The temperature of the split/splitless injector, transfer line, and ion source were set to 250°C. The oven temperature gradually increased from an initial temperature of 70°C to a final temperature of 350°C at a rate of 9°C/min. To prevent solvent contamination, a solvent delay of 340 seconds was implemented. The metabolites were ionised and fractionated using a 70 eV ion pulse. Mass spectra, spanning an m/z range of 35–800, were captured at a frequency of 20 scans per second. The collected data, including chromatograms and mass spectra, were evaluated and interpreted using ChromaTOF 4.7 and TagFinder 4.1 software (Luedemann et al., 2008), and the Golm Metabolome Database (Kopka et al., 2005).For details see Supplemental Data S1

**Mass spectrometry was used for proteome analysis.**

The total leaf material of five-week-old plants was analyzed in three independent biological replicates. Protein extraction and trypsin digestion were performed following the protocol described in Marino et al. (2019). Liquid chromatography-tandem mass spectrometry (LC-MS/MS) was performed as previously described, but peptides were separated over a 90-minute linear gradient of 5-80% (v/v) ACN, according to (Espinoza-Corral et al., 2023). Raw files were processed using MaxQuant software version 2.1.0.0 (Cox and Mann, 2008). The peak lists were searched against the Arabidopsis reference proteome (Uniprot, www.uniprot.org) using default settings with 'mach-between-runs' enabled. The proteins were quantified using the label-free quantification algorithm (LFQ) (Cox et al., 2014). The downstream analysis was performed using Perseus version 2.0.6.0 (Tyanova et al., 2016). For further analysis, potential contaminants, proteins identified only by site modification, and reverse hits were excluded. The protein groups that could be quantified by the LFQ algorithm in at least two of the three replicates in at least one condition were included. The LFQ intensities were $log_2$-transformed, and missing values were imputed from a normal distribution using standard settings in Perseus. For more information, refer to Supplemental Data S2.

**For transcriptome analyses,**

total leaf material from four-week-old plants was used in three biological replicates. Plants were grown as described above, and samples were harvested either after 3 hours of light (ML, control) or after three hours of high light (HL) at 1000µmol photons m−2 s−1. Novogene (Beijing, China) performed RNA library preparation and transcriptome sequencing. To generate the cDNA libraries, poly(A)+ RNA enrichment and mRNA fragmentation were performed, followed by random-prime cDNA synthesis to remove ribosomal RNA. The reads were captured using Illumina sequencing (NovaSeq 6000) and were paired-end sequenced with a length of 150 base pairs. To assess the quality of the reads, we used the bioinformatics program Hisat2 to map them to the Arabidopsis reference genome. Novogene conducted further bioinformatic analysis. We identified differentially expressed genes (DEGs) using DESeq2 analysis (Love et al., 2014) with an adjusted p-value threshold of <0.05.


**Cox J, Hein MY, Luber CA, Paron I, Nagaraj N, Mann M** (2014) Accurate proteome-wide label-free quantification by delayed normalization and maximal peptide ratio extraction, termed MaxLFQ. Mol Cell Proteomics **13:** 2513-2526

**Cox J, Mann M** (2008) MaxQuant enables high peptide identification rates, individualized p.p.b.-range mass accuracies and proteome-wide protein quantification. Nat Biotechnol **26:** 1367-1372

**Espinoza-Corral R, Schwenkert S, Schneider A** (2023) Characterization of the preferred cation cofactors of chloroplast protein kinases in Arabidopsis thaliana. FEBS Open Bio **13:** 511-518

**Garcia-Molina A, Kleine T, Schneider K, Muhlhaus T, Lehmann M, Leister D** (2020) Translational Components Contribute to Acclimation Responses to High Light, Heat, and Cold in Arabidopsis. iScience **23:** 101331

**Kopka J, Schauer N, Krueger S, Birkemeyer C, Usadel B, Bergmuller E, Dormann P, Weckwerth W, Gibon Y, Stitt M, Willmitzer L, Fernie AR, Steinhauser D** (2005): the Golm Metabolome Database. Bioinformatics **21:** 1635-1638

**Love MI, Huber W, Anders S** (2014) Moderated estimation of fold change and dispersion for RNA-seq data with DESeq2. Genome Biol **15:** 550

**Stitt M, McC.Lilley R, Gerhardt R, Heldt HW** (1989) Metabolite levels in specific cells and subcellular compartments of plant leaves. Meth. Enzymol **174:** 518-552

**Straube H, Niehaus M, Zwittian S, Witte CP, Herde M** (2021) Enhanced nucleotide analysis enables the quantification of deoxynucleotides in plants and algae revealing connections between nucleoside and deoxynucleoside metabolism. Plant Cell **33:** 270-289

**Tyanova S, Temu T, Sinitcyn P, Carlson A, Hein MY, Geiger T, Mann M, Cox J** (2016) The Perseus computational platform for comprehensive analysis of (prote)omics data. Nat Methods **13:** 731-740
