## Supplemental Table 1 for "UPP affects chloroplast development by interfering with chloroplast proteostasis"

Supplemental Table S1. Primers used in this study

| Name | Sequence | Purpose |
|---|---|---|
| *ami-upp-3, -4* and *-5* | | |
| UPP1_I miR-s | gaTATACATGAGTAATCTCGCTAtctctcttttgtattcc | |
| UPP1_II miR-a | gaTAGCGAGATTACTCATGTATAtcaaagagaatcaatga | Oligos for amiRNA "TATACATGAGTAATCTCGCTA" |
| UPP1_III miR*s | gaTAACGAGATTACTGATGTATTtcacaggtcgtgatatg | |
| UPP1_IV miR*a | gaAATACATCAGTAATCTCGTTAtctacatatatattcct | |
| *ami-upp-1, -2, -6, -7* and *-8* | | |
| UPP3_I miR-s | gaCTGCGGTTGTTCCGATATTAAtcaaagagaatcaatga | |
| UPP3_II miR-a | gaCTGCGGTTGTTCCGATATTAAtcaaagagaatcaatga | Oligos for amiRNA "TTAATATCGGAACAACCGCAG" |
| UPP3_III miR*s | gaCTACGGTTGTTCCCATATTATtcacaggtcgtgatatg | |
| UPP3_IV miR*a | gaATAATATGGGAACAACCGTAGtctacatatatattcct | |
| UPP_RT_fwd | ttgcggaacacgcatcctca | |
| UPP_RT_rev | gtttctgtcccgaaactgcg | qRT-PCR primer - UPP transcript |
